# The *Capicua* C1 Domain is Required for Full Activity of the CIC::DUX4 Fusion Oncoprotein

**DOI:** 10.1101/2024.06.06.597815

**Authors:** Cuyler Luck, Kyle A. Jacobs, Ross A. Okimoto

## Abstract

Rearrangements between genes can yield neomorphic fusions that drive oncogenesis. Fusion oncogenes are made up of fractional segments of the partner genes that comprise them, with each partner potentially contributing some of its own function to the nascent fusion oncoprotein. Clinically, fusion oncoproteins driving one diagnostic entity are typically clustered into a single molecular subset and are often treated a similar fashion. However, knowledge of where specific fusion breakpoints occur in partner genes, and the resulting retention of functional domains in the fusion, is an important determinant of fusion oncoprotein activity and may differ between patients. This study investigates this phenomena through the example of CIC::DUX4, a fusion between the transcriptional repressor capicua (*CIC*) and the double homeobox 4 gene (*DUX4*), which drives an aggressive subset of undifferentiated round cell sarcoma. Using a harmonized dataset of over 100 patient fusion breakpoints from the literature, we show that most bona fide CIC::DUX4 fusions retain the C1 domain, which is known to contribute to DNA binding by wild type CIC. Mechanistically, deletion or mutation of the C1 domain reduces, but does not eliminate, activation of *CIC* target genes by CIC::DUX4. We also find that expression of C1-deleted CIC::DUX4 is capable of exerting intermediate transformation-related phenotypes compared with those imparted by full-length CIC::DUX4, but was not sufficient for tumorigenesis in a subcutaneous mouse model. In summary, our results suggest a supercharging role for the C1 domain in the activity of CIC::DUX4.

**Significance Statement:** Fusion oncogenes are neomorphic entities comprised of fractional coding sequences from two partner genes that have been inappropriately rearranged. The functional domains contributed by the partner genes shape the function of the resulting fusion. We use CIC::DUX4, a transcription factor fusion that drives an ultra-rare soft tissue sarcoma, to explore how preferential retention of a partner gene domain may influence the activity of the overall fusion. Our results indicate that the capicua C1 domain is retained in most CIC::DUX4 transcripts and is required for full activity of the CIC::DUX4 oncoprotein. We demonstrate that knowledge of where breakpoints occur and the resulting impact this has on the retention of functional domains can teach us about fusion behavior.

## Introduction

When DNA replication and repair pathways fail to faithfully process genomic DNA, one possible outcome is the formation of fusion genes comprised of portions of two individual partner genes that have been rearranged into a single gene. In the case of many cancer types, such events give rise to fusion oncogenes, which are transcribed and translated into fusion oncoproteins that deviate from the behavior of their wild-type partners. Transcription factor (TF) fusions represent a subset of fusion oncogenes that drive multiple tumor types, including sarcomas, through chromatin remodeling, enhancer hijacking, and target gene dysregulation (1–6). In such tumor types, the TF fusion is often one of few, if not the sole, genetic alterations present (7, 8), suggesting they are key driver events.

While fusion oncoproteins are neomorphic entities not observed in healthy cells, the functional domains guiding their form, function, and regulation are inherited from the normal partner genes that comprise them. This can directly determine how the fusion executes its oncogenic function: for example, PAX3::FOXO1 (fusion-positive rhabdomyosarcoma) binds DNA at sites determined by the *PAX3* DNA binding domains, while *FOXO1* provides transactivating capacity (9, 10). Beyond directly shaping output, partner gene domains can also determine regulation of the fusion, as is the case for ERK-mediated degradation of CIC::DUX4 (CIC-rearranged sarcoma) due to retention of the *CIC* ERK-binding domain in the fusion oncoprotein (11, 12). Due to the critical roles they play in the activity of the overall fusion, knowing which functional domains are retained from partner genes can be informative in understanding how a fusion acts or can be acted upon. Importantly, fusion breakpoints and transcripts can be variable between patients with the same tumor types (13–15), which may result in different exons and potentially variable functional domains being retained in any individual fusion. Despite these possible molecular differences, it is often assumed that all fusion oncoproteins made up of a given set of partner genes operate in a similar fashion.

A particularly interesting class of TF fusions are those involving rearrangements of capicua (*CIC*) with one of several 3’ partner genes, typically *DUX4* (but also including *NUTM1*, *LEUTX*, and *FOXO4*). Wild type (WT) *CIC* is a transcriptional repressor discovered in *Drosophila* which is involved in developmental processes (16, 17) and acts as a tumor and metastasis suppressor predominantly through the repression of *PEA3 (ETV1, ETV4, ETV5)* family members and negative MAPK-pathway (*DUSP4/6)* regulators (18–21). As a tumor suppressor, WT *CIC* is genetically deleted, mutated, or otherwise functionally inactivated in several types of cancers as we have previously reviewed (22). Interestingly, when fused to *DUX4*, CIC::DUX4 becomes an oncogenic neomorphic transcriptional activator of CIC target genes (6, 23–25) and drives undifferentiated round cell sarcoma with a dismal prognosis (26) and limited treatment options. This contrast between WT CIC acting as a tumor suppressor and CIC::DUX4 being an oncogene provides a rare opportunity to take knowledge learned from natural evolution of CIC-inactivation in WT CIC cancers and apply those strategies to potentially disable the CIC::DUX4 fusion oncoprotein.

We have taken a particular interest in the C1 domain of *CIC*, a C-terminal DNA binding domain that cooperates with the HMG box to engage DNA at target sites, as has been best shown by Forés and colleagues among others (18, 27, 28). Their work highlighted that the C1 domain is a mutational hotspot in glioma and showed that in *Drosophila*, truncation of the C1 domain in a synthetic fly-human CIC::DUX4 construct partially rescues phenotypes imparted by expression of the fully intact fusion. However, the role of the C1 domain in CIC::DUX4 activity has not been formally tested in a human or mammalian context. The C1 domain is retained in the fusion transcripts of the few existing patient-derived CIC::DUX4 sarcoma cell lines (6, 29–31), but there has been no large-scale comprehensive analysis of CIC::DUX4 patient fusion breakpoints and which functional domains are retained in their fusion oncoproteins. This limits our understanding of how frequently the C1 domain is included in CIC::DUX4 oncoproteins in patients, making generalizations to patient biology difficult.

In this study we leverage the first harmonized database of CIC-rearranged breakpoints from patients along with cellular models to interrogate the role of the C1 domain in mammalian CIC::DUX4 activity. We find that the C1 domain is retained in the majority of bona-fide CIC::DUX4 patient fusions, with the *CIC* breakpoint typically constrained to a relatively small C-terminal region. Deletion or mutation of the C1 domain attenuated, but did not entirely eliminate, activation of target genes in multiple cell models with ectopic expression of CIC::DUX4. This translated to phenotypic outcomes, where C1-deleted CIC::DUX4 was capable of driving intermediate phenotypes related to blocking differentiation and contact-inhibited growth. However, expression of C1-deleted CIC::DUX4 was not capable of forming tumors in an *in vivo* subcutaneous implantation model, in contrast to expression of the full-length CIC::DUX4 fusion oncoprotein. Taken together, these results suggest that the C1 domain is necessary for maximal activity of the CIC::DUX4 fusion oncoprotein in mammalian systems.

## Results

### The Majority of Bona Fide CIC::DUX4 Fusions in Patient Tumors Retain the C1 Domain

We first hypothesized that if the C1 domain is functionally important for human sarcomagenesis, then the *CIC* breakpoints observed in CIC-fusion transcripts would likely non-randomly occur after the 3’ end of the C1 domain, preserving it in the resulting fusion oncoprotein. To our knowledge, no comprehensive databases of CIC-fusion breakpoints exist, and reports describing CIC-fusion transcripts often use different reference sequences, complicating analysis. To overcome this challenge, we trawled the literature for reports of any CIC-rearranged fusion transcripts from patients, manually aligning all collected breakpoints to reference sequences (Figure 1A). We were able to identify 108 nucleotide-level breakpoints from 41 different publications for four different *CIC* fusions (*DUX4*, *NUTM1*, *LEUTX*, and *FOXO4*, Figure 1B), with an additional 26 exon-level breakpoints (full dataset available in Supplemental Dataset S1). For further detailed analysis we focused on breakpoints derived from RNA data where information on both partners were available. Interestingly, while *CIC* breakpoints for *DUX4*, *LEUTX*, and *FOXO4* fusions were largely at the far C-terminal end of the gene, those for *NUTM1* fusions were often relatively earlier in the coding sequence, including upstream of the C1 domain (Supplemental Figure S1A). However, low sample numbers for the exceptionally rare non-*DUX4* fusions limited the generalizations that could be drawn for *LEUTX, FOXO4, and NUTM1* fusions.

**Figure 1.**
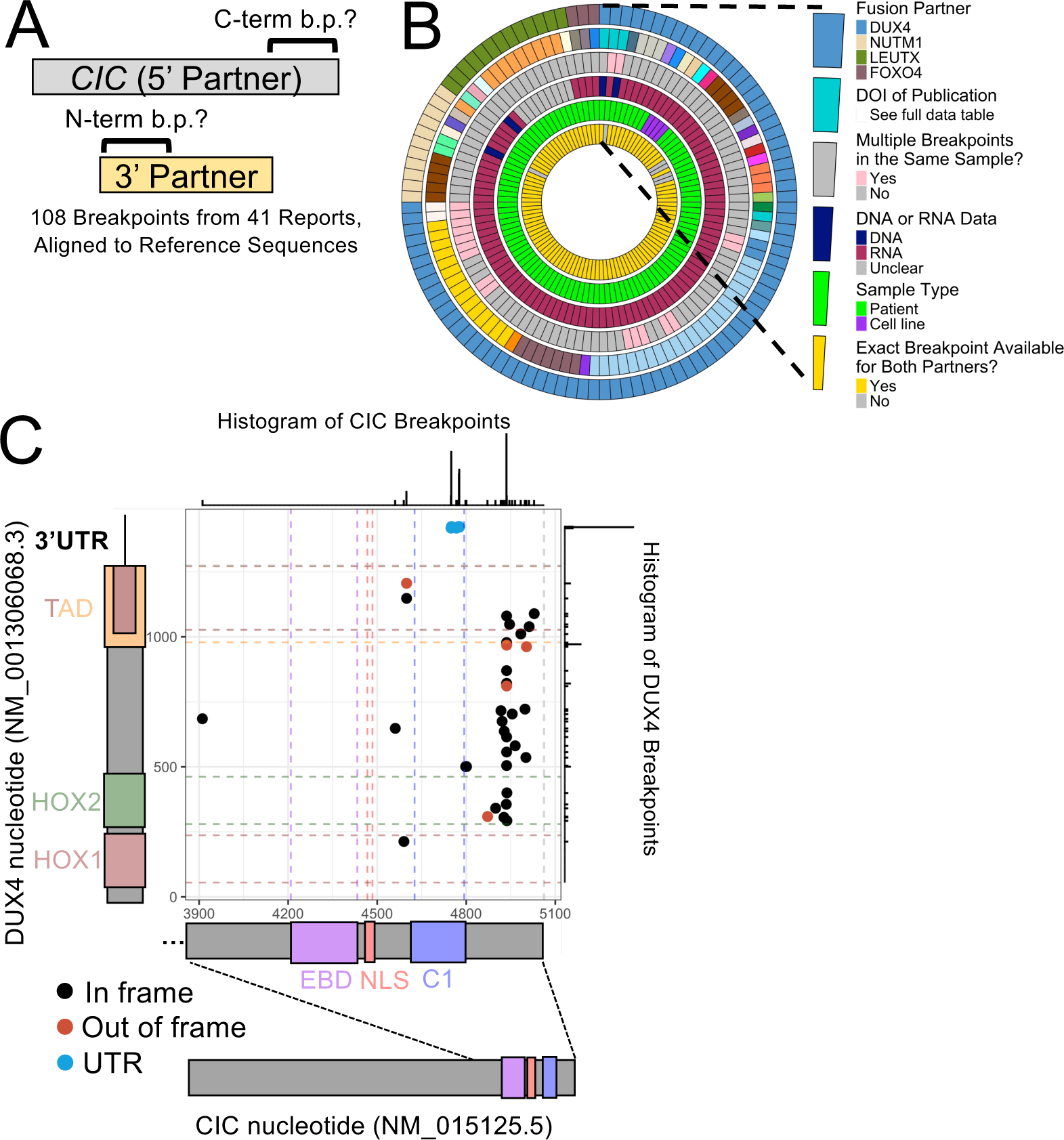
The capicua C1 domain is retained in most bona fide CIC::DUX4 fusion transcripts. (A) Schematic describing harmonization of patient breakpoint (b.p.) data. (B) Circos plot of all 108 nucleotide-level breakpoints gathered for CIC-fusion genes. (C) Scatterplot of breakpoint coordinates from 74 CIC::DUX4 transcripts where the data source was RNA and both partner breakpoints were available. Each data point is one breakpoint, histograms are intended to aid with overplotting. EBD: ERK-binding domain, NLS: nuclear localization signal, C1: C1 domain, HOX1/2: homeodomains 1/2, TAD: two overlapping putative transactivation regions identified in *DUX4*.

Since CIC::DUX4 fusions had the most breakpoints to analyze, we next focused on the 74 nucleotide-level CIC::DUX4 breakpoints which met the aforementioned selection criteria. Interestingly, we observed a large number of *CIC* breakpoints in CIC::DUX4 fusions that occurred within the C1 domain (n=32 of 74, 43%), which we would expect to disrupt C1 domain function (Supplemental Figure S1A). However, closer analysis of the *DUX4* breakpoint in these samples indicated that these fusions are actually between *CIC* and the 3’UTR of *DUX4*, which is consistent with prior observations of these cases in their primary reports (32–36) (Figure 1C). These “*CIC*::UTR” fusions have been described to often have STOP codons shortly after the breakpoint, leading to a truncated CIC protein lacking a C-terminal region infringing on the C1 domain, which has been hypothesized to explain their apparent tumorigenic nature (32). To model this phenomenon, we used an HA-CIC::DUX4 plasmid (6) to clone one HA-CIC::UTR fusion based on case number 4 from Cocchi and colleagues (35) (Supplemental Figure S1B). We found that while its expression in 293T cells significantly increased transcription of some known CIC target genes compared with the negative control, the magnitude of change in gene expression was exceptionally small compared with that observed following transfection of a full-length HA-CIC::DUX4 (Supplemental Figure S1C-D).

Of the remaining 42 CIC::DUX4 breakpoints, we determined the majority to be genuine in-frame fusions (n=37 of 42, 88%). Almost all of these bona fide CIC::DUX4 fusions retained the C1 domain, with the *CIC* breakpoint occurring after the 3’ end of the C1 domain in 32 out of 37 bona fide CIC::DUX4 breakpoints (86%, Figure 1C). This is exceptionally constrained compared to the distribution of *DUX4* breakpoints in the same samples, where the *DUX4* breakpoints are observed to occur almost anywhere between the HOX1 DNA-binding domain and the C-terminal region previously described as important for recruitment of p300 and transactivation capacity (37, 38) (Figure 1C). Given that the extremely narrow C-terminal window where most *CIC* breakpoints occur in bona fide CIC::DUX4 fusions results in retention of the C1 domain in the fusion oncoprotein, we interpret this to suggest a likely functional contribution of the C1 domain to the activity of the fusion in a human context.

### Deletion or Mutation of the C1 Domain Partially, but not Completely, Impairs Target Gene Induction by CIC::DUX4 in 293T Cells

In order to test the functional impact of the C1 domain on human CIC::DUX4 activity, we deleted the C1 domain, the HMG box, or both from an established HA-CIC::DUX4 expression construct (6) and expressed them in 293T cells (Figure 2A). Since the HMG box and the C1 domain have been shown to be required together for proper binding of CIC target DNA sites (27, 28, 39), the HMG/C1 co-deletion mutant functions as a DNA-binding-dead CIC::DUX4 negative control. All constructs expressed at the expected protein level (Figure 2B). Intriguingly, while deletion of the C1 domain significantly impaired transcriptional induction of CIC target genes, the observed activation remained higher than the DNA-binding-dead control in the majority of tested targets (Figure 2C, Supplemental Figure S2A). To test if we could replicate these findings without large truncating deletions in the same functional domains, we engineered point mutations at residues R201 (HMG box, mutated to W) and R1515 (C1 domain, mutated to H) based on previously described hotspot mutations found in CIC-mutated gliomas (27) (Figure 2A). Indeed, we again observed that a point mutation in the C1 domain significantly reduced, but did not eliminate, induction of CIC target genes (Figure 2D and 2E, Supplemental Figure S2B). We orthogonally validated these results by testing activation of an *ETV5* promoter-luciferase construct carrying the consensus CIC binding sequence by the deletion mutants, finding again that the C1-deleted mutant showed incomplete abrogation of transcriptional activation (Supplemental Figure S2C). Immunofluorescence microscopy of full-length or deletion-mutant HA-CIC::DUX4 transfected cells indicated that the deletions were not overtly impacting nuclear localization (Figure 2F). Together, these results indicate that the C1 domain of CIC::DUX4 enhances activation of CIC::DUX4 target genes in 293T cells.

**Figure 2.**
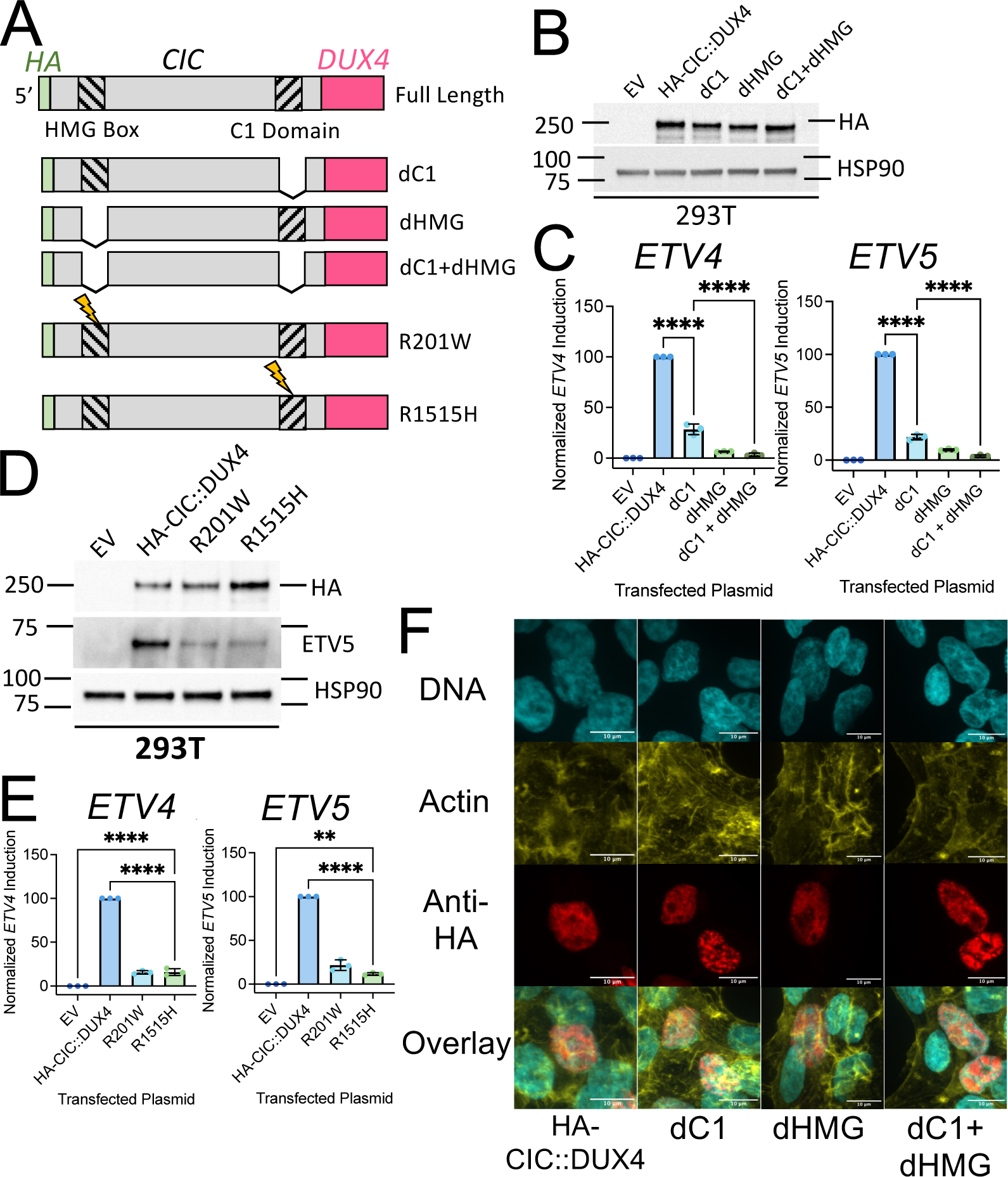
Deletion of the C1 domain reduces, but does not eliminate, CIC-target gene induction without overtly impacting CIC::DUX4 protein expression or localization. (A) Schematic of the engineered CIC::DUX4 deletion or point mutation variants. (B) Immunoblot of 293T cells approximately 48 hours after transfection with empty vector (EV) or the indicated constructs, data representative of three independent experiments. (C) Normalized RT-qPCR measurement of target gene (*ETV4* and *ETV5*) induction in 293T cells approximately 48 hours after transfection with EV or the indicated constructs. Each data point represents the mean of one of three independent experiments, error bars indicate standard deviation, **** = p < 0.0001 by one-way ANOVA and Šidák’s multiple comparisons test. (D) Immunoblot of 293T cells approximately 48 hours after transfection with empty vector (EV) or the indicated constructs, data representative of three independent experiments. (E) Normalized RT-qPCR measurement of target gene (*ETV4* and *ETV5*) induction in 293T cells approximately 48 hours after transfection with EV or the indicated constructs. Each data point represents the mean of one of three independent experiments, error bars indicate standard deviation, **** = p < 0.0001 and ** = p < 0.01 by one-way ANOVA and Šidák’s multiple comparisons test. (F) Confocal immunofluorescence imaging of 293T cells approximately 48 hours after transfection with the labeled constructs. DNA visualized with DAPI, actin visualized with rhodamine-phalloidin, 60x objective used for imaging, scale bars indicate 10 μm, representative cells chosen from one experiment.

### Stable NIH/3T3 and C2C12 Clones Validate Differential Induction of ETV5 by C1-Intact or C1-Deleted CIC::DUX4

To broaden our findings beyond transient expression systems and 293T cells, we next aimed to stably express CIC::DUX4 or the C1-deleted mutant in cell lines more reflective of sarcoma biology. We chose to employ mouse NIH/3T3 fibroblasts and mouse C2C12 myoblasts, as both have previously been used to study CIC::DUX4 and other sarcomas (6, 24, 25) and represent the putative progenitor of soft tissue sarcomas (mesenchymal lineage - myoblast). We used a combination of retroviral transduction and fluorescence-activated cell sorting to generate clonal cell lines expressing empty vector (“EV”), full-length CIC::DUX4 (“CD4”), or C1-deleted CIC::DUX4 (“dC1”) with an IRES-EGFP reporter (Figure 3A). All but one evaluated NIH/3T3 clones expressed the proper transgenes (Supplemental Figure S3 A and B). Interestingly, while the C2C12 EV and dC1 clones all expressed the proper transgenes, only one of six tested C2C12 CD4 clones displayed measurable full-length CIC::DUX4 protein expression (Supplemental Figure S3C). Across both NIH/3T3 and C2C12 clones, dC1 clones consistently expressed higher protein levels of mutant CIC::DUX4 than CD4 clones expressed full-length CIC::DUX4, which we speculate is in part due to toxicity and intolerance of high full-length fusion expression as previously described with other fusion oncoproteins (40). NIH/3T3 clones expressing either CIC::DUX4 version expressed less EGFP than the EV clones, suggesting that CIC::DUX4 transgenes were transcribed less than EV transgenes. Despite having lower fusion oncoprotein expression, CD4 clones expressing full-length CIC::DUX4 demonstrated higher levels of ETV5 protein expression than dC1 clones expressing C1-deleted CIC::DUX4, corroborating our findings from 293T cells.

**Figure 3.**
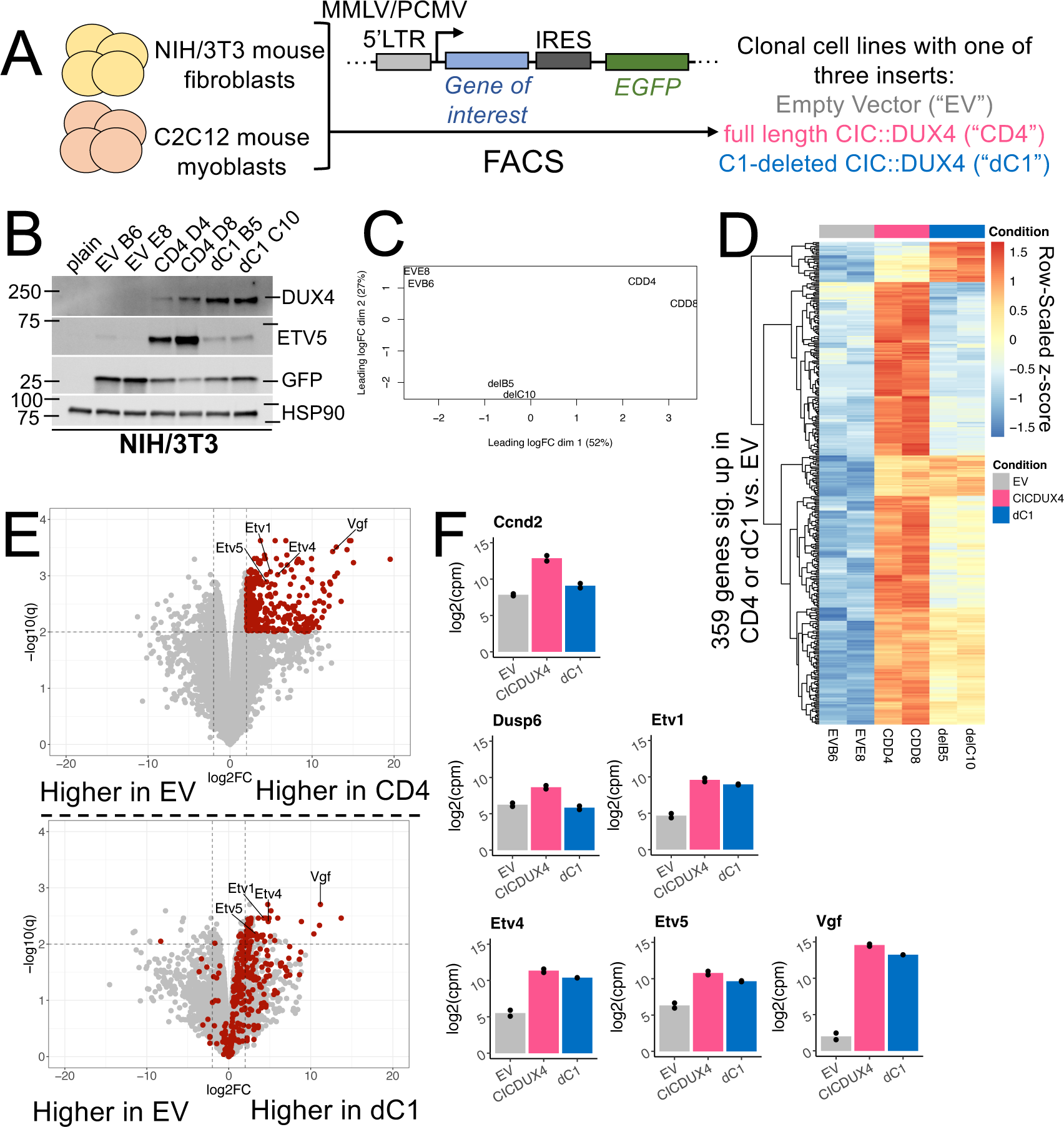
Stable expression of C1-deleted CIC::DUX4 in clonal NIH/3T3 cells drives a largely attenuated transcriptomic program defined by full-length CIC::DUX4 expression. (A) Schematic describing how clonal populations of transduced NIH/3T3 and C2C12 cells were generated. (B) Immunoblot of selected clonal NIH/3T3 cell lines in comparison with plain untransduced NIH/3T3. Clones were evaluated for transgene expression three independent times including the initial clone screening. (C) Multi-dimensional scaling plot of NIH/3T3 clone RNA-seq data after processing with edgeR. (D) Row-scaled heatmap of 359 significantly upregulated (log_2_ fold change > 2, q < 0.01) genes in either of the CD4 vs EV or dC1 vs EV comparisons across six NIH/3T3 clones. (E) Volcano plots of differentially expressed genes between full-length CIC::DUX4 (CD4) or C1-deleted CIC::DUX4 (dC1) clones and empty vector (EV) clones. Maroon-colored genes were significantly upregulated (log_2_ fold change > 2, q < 0.01) in the CD4 vs EV comparison. Select known or high-confidence CIC/CIC::DUX4 target genes are labeled. (F) log_2_(counts per million) measurements for six selected known or high-confidence CIC/CIC::DUX4 target genes in NIH/3T3 clones, grouped by transduction. Bars represent mean values. In the edgeR differential expression analysis using quasi-likelihood F tests, all six genes were significantly different for CD4 vs EV (q < 0.01) and only four genes (*Etv1*, *Etv4*, *Etv5*, *Vgf*) were significantly different (q < 0.01) for dC1 vs EV.

### Deletion of the C1 Domain Impairs, but Does Not Fully Eliminate, the Transcriptional Program Controlled by CIC::DUX4 in NIH/3T3 Cells

We selected two clones each from the NIH/3T3 stable cell lines to continue with unbiased transcriptional analysis via bulk RNA-sequencing (Figure 3B). Treating the two clones within each group as biological replicates, multidimensional scaling indicated that the C1-deleted clones had a transcriptional profile in-between those of either the EV or full-length CIC::DUX4 clones (Figure 3C). A heatmap of all genes significantly upregulated (log_2_ fold change > 2, q < 0.01) in either the CD4 or dC1 conditions vs. EV revealed distinct blocks of genes regulated differently by the two versions of CIC::DUX4 (Figure 3D). Most genes were strongly upregulated by full-length CIC::DUX4 but only weakly activated or not activated at all by C1-deleted CIC::DUX4. However, there was a small block of genes whose activation did not depend on which CIC::DUX4 variant was expressed, and a subset of genes that were more strongly upregulated by C1-deleted CIC::DUX4 when compared to full length CIC::DUX4. Volcano plots of either CD4 or dC1 clones compared with EV clones revealed that expression of essentially all genes significantly activated by full-length CIC::DUX4 was attenuated in the C1-deleted clones (Figure 3E). Focusing on six validated or high-confidence CIC::DUX4 target genes, we observed that while all six targets had lower expression in the dC1 clones compared to the CD4 clones, even this attenuated expression was still significantly higher than the EV clones for *Etv1*, *Etv4*, *Etv5*, and *Vgf* (Figure 3F). While loss of the C1 domain impaired the CIC::DUX4 mediated transcriptional program in stable NIH/3T3 clones, it remained unclear whether this would translate to differences in functional phenotypes driven by these fusion oncoproteins.

### Deletion of the C1 Domain Yields Intermediate CIC::DUX4 Driven, Transformation-Related Phenotypes

Co-opting differentiation pathways is a known hallmark of some fusion oncoproteins, which to our knowledge has not been defined for CIC::DUX4. Thus, we first asked if CIC::DUX4 +/- the C1 domain can interfere with C2C12 differentiation *in vitro*. C2C12 cells plated at confluency and serum-starved differentiate into myotubes, in the process expressing key TF markers including myogenin (*MYOG*) (41). We subjected three of our C2C12 clonal cell lines, one each of EV, CD4, and dC1, to such an assay and used myogenin expression as a proxy for myotube differentiation. CIC::DUX4-expressing clones demonstrated reduced myogenin protein expression relative to the EV clone, with the full-length CIC::DUX4 clone having no detectable protein expression and the C1-deleted clone having minimal expression at days 4 and 6 (Figure 4A). To test that the observed change in myogenin expression was a direct effect of CIC::DUX4 transgene expression we next performed a rescue experiment with the same differentiation assay where an siRNA pool against human *CIC* was used to deplete CIC::DUX4 expression during differentiation. siRNA targeting human *CIC* had no detectable effect on myogenin expression in the EV clone compared with control, while we again observed that control-treated CIC::DUX4 expressing clones yielded no (full length CIC::DUX4) or strongly reduced (C1-deleted CIC::DUX4) myogenin (Figure 4B). Importantly, this was partially reversible with siCIC treatment and accompanying mild knockdown of CIC::DUX4 protein at day 5 (Figure 4B). Taken together, these results suggest that CIC::DUX4 expression does block myotube differentiation of C2C12 cells *in vitro*, and that C1-deleted CIC::DUX4 is capable of driving a slightly attenuated but similar phenotype.

**Figure 4.**
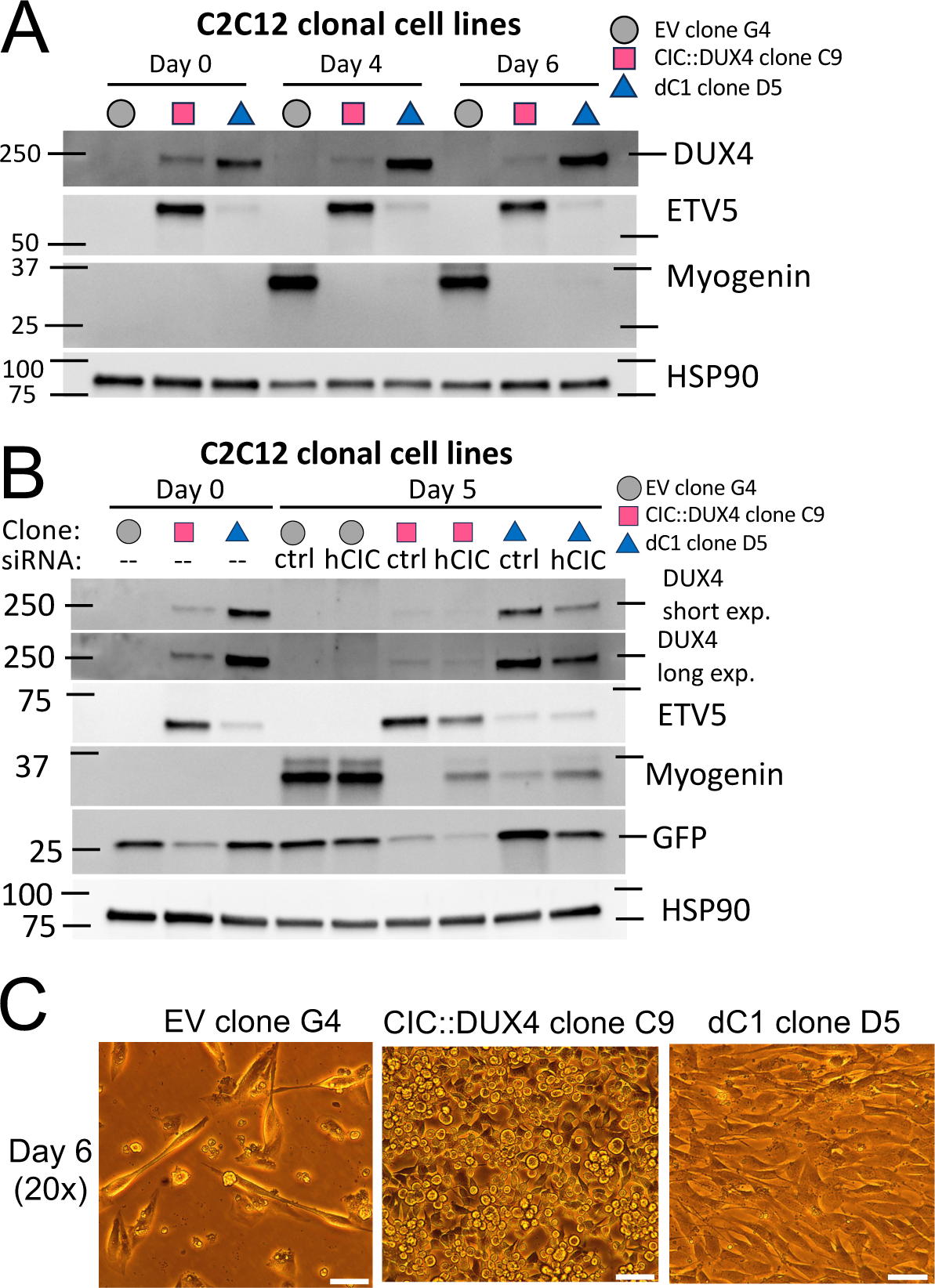
Expression of CIC::DUX4 with or without the C1 domain can block differentiation and alter the growth patterns of clonal C2C12 cells. (A) Immunoblot of clonal C2C12 cells after differentiation for the indicated times, representative of two independent experiments. (B) Immunoblot of clonal C2C12 cells after differentiation for the indicated times and following transfection with non-targeting (ctrl) or human *CIC* targeting (hCIC) siRNA. Representative of two independent experiments. (C) Microscopy images of C2C12 clones after six days of differentiation. Imaged using a 20x objective, scale bar indicates 0.05 mm, representative of two independent experiments.

During the process of the initial differentiation experiments, we recorded images of differentiating cells at several time points with the intention of visualizing elongation of cells into myotubes. While we did observe this in the EV clone, we unexpectedly observed phenotypic differences in cell growth and contact inhibition. By day 4, the EV clone had clearly begun to either die off or start elongating into myotubes, which continued into day 6 (Figure 4C, Supplemental Figure S4A). In contrast, the full-length CIC::DUX4 clone continued proliferating throughout the entire experiment, appearing to grow vertically on top of its initial monolayer (Figure 4C, Supplemental Figure S4A). Strikingly, while the C1-deleted clone was also capable of continued proliferation after switching to differentiation media, it remained constrained to a single monolayer, in contrast to the full-length CIC::DUX4 clone (Figure 4C, Supplemental Figure S4A).

As an additional means of validating these cell-cell contact and growth phenotypes, we leveraged our NIH/3T3 clones to test 3D growth phenotypes in a hanging drop assay. While the EV clones were capable of forming small tightly packed spheroids with few isolated cells, the full-length CIC::DUX4 clones displayed atypical growth patterns with many free-floating dissociated cells not associated with larger assemblies (Supplemental Figure S5). As with other experiments, the C1-deleted clones demonstrated an intermediate phenotype, with one clone (B5) forming fairly EV-like spheroids while the other (C10) formed small spheroids that clumped up in chains or larger structures (Supplemental Figure S5). Taken together, these results suggest that deletion of the C1 domain yields phenotypes intermediate to those conferred by full-length CIC::DUX4, including some transformation-related phenomena such as differentiation-blocking and non-contact-inhibited growth.

### Expression of C1-Deleted CIC::DUX4 in C2C12 Clones is not Sufficient to Drive Tumorigenesis in Nude Mice

Since we consistently observed intermediate activity of the C1-deleted CIC::DUX4 compared with full-length CIC::DUX4 across several *in vitro* assays, we next aimed to test the impact on tumor growth *in vivo*. We subcutaneously implanted our C2C12 clones into the flanks of immunodeficient (nude) mice (Figure 5A). C2C12 myoblasts are representative of a mesenchymal stem cell progenitor and they only generate lesions over fairly long timeframes when implanted in mice (42–44). Consistent with these findings, only half of the injection sites (5/10, 50%) of EV-injected mice developed small (less than 14 mm^3^) slow-growing lesions over the course of the 30 day experiment (Figure 5B, Supplemental Figure S6A). In contrast, full length CIC::DUX4-injected mice rapidly developed tumors (10/10, 100%) of at least 50 mm^3^ by day 14 post-injection (Figure 5B, Supplemental Figure S6A). Three CD4-injected mice had to be sacrificed at day 14 due to ulcerating tumors, while the remaining two CD4-injected mice were sacrificed at day 16 to avoid ulceration from their established tumors. Unexpectedly, mice injected with C1-deleted CIC::DUX4 expressing cells did not develop measurable tumors at study termination, with only one injection site having a small lesion of 6 mm^3^ (1/10, 10%, Figure 5B, Supplemental Figure S6A). Western blot analysis of explanted tissue could not reliably detect CIC::DUX4 protein expression in any tumor sample but did reveal increased ETV5 levels in full-length CIC::DUX4 tumors compared with EV or dC1 tissue (Figure 5C). We were able to confirm CIC::DUX4 transcript expression at the RNA level for full length CIC::DUX4 tumors (Figure 5D) but did not have remaining tissue to evaluate dC1 lesions. These results suggest that in this particular model, C1-deleted CIC::DUX4 was not sufficient to drive tumorigenesis.

**Figure 5.**
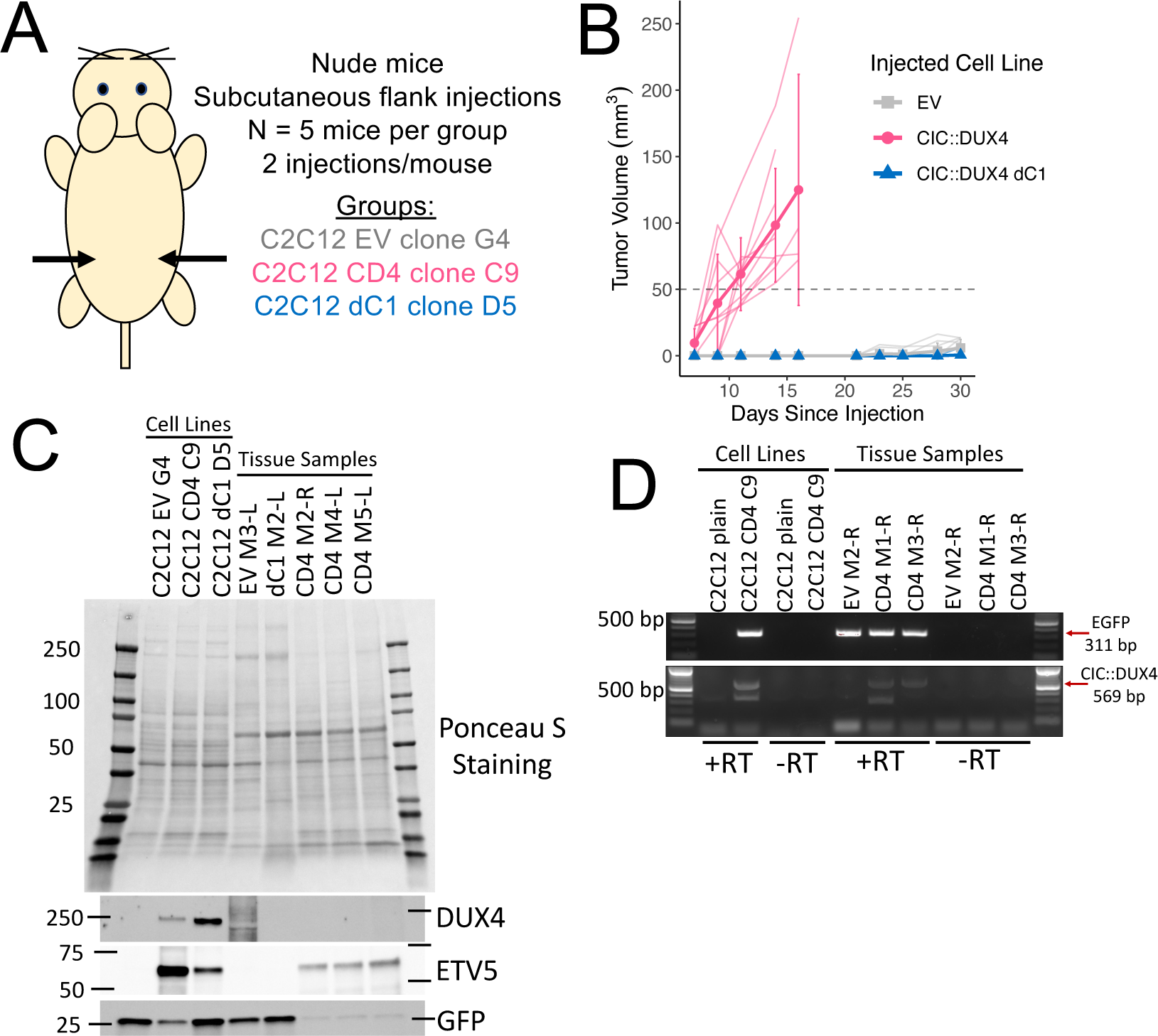
C1-deleted CIC::DUX4 expressing C2C12 cells do not initiate tumor formation in nude mice. (A) Schematic of clonal C2C12 subcutaneous implantation experimental design. (B) Plot of tumor volumes versus time, thin transparent lines indicate individual injection sites while thick opaque lines and data points represent averages. Error bars represent standard deviation. Note that three full-length CIC::DUX4 injected mice were sacrificed at day 14, and the remaining two mice in that group were sacrificed at day 16. (C) Ponceau S staining and immunoblots of clonal C2C12 cell lines or explanted tumor tissue. Tumor tissue is labeled by injected cell line, mouse number, and flank side. Protein analysis was performed once due to inability to detect CIC::DUX4 expression in tissue. (D) Agarose gel electrophoresis of PCR products from cDNA derived from C2C12 cell lines or explanted tumor tissue. Arrows indicate expected sizes of positive PCR products for the indicated transcripts. The smaller band at approximately 300-350 bp in the CIC::DUX4 reaction is nonspecific. +/-RT refers to inclusion/exclusion of reverse transcriptase in the cDNA generation step. Red pixels within bands indicates saturated signal. The same cDNA from tissue samples was tested two separate times with the same results.

## Discussion

Fusion oncogenes, including TF fusions, are often defined at the gene level (i.e. *EWS*-*FLI1*, not *EWSR1* exon 7 – *FLI* exon 5). This is certainly understandable given the vast number of fusion oncogenes driving human disease and the requirement for rapid and precise clinical and diagnostic interpretation. However, specific breakpoints where partner genes are fused in the context of these rearrangements are not always identical (13–15). We therefore argue that there is value in understanding where those sites occur and how the inclusion or exclusion of key regulatory domains from the partner genes impact fusion oncoprotein function.

Our analysis of CIC::DUX4 breakpoints described in the literature reveals that in bona fide CIC::DUX4 fusion transcripts, *DUX4* breakpoints occur almost anywhere after the HOX1 domain and prior to the C-terminal region responsible for transactivation and interaction with p300. This distribution of *DUX4* breakpoint locations is consistent with the current model that DUX4 contributes its p300-recruitment and transactivation capacities to the CIC::DUX4 fusion oncoprotein, but not its DNA-binding specificity. In contrast to the variability observed for *DUX4* breakpoints, *CIC* breakpoints occur almost exclusively at the very 3’ end of the coding sequence, retaining essentially all of the *CIC* transcript (4500+ nucleotides) in the fusion. While this could be due to factors like sequence-specific properties which make those regions more likely to form translocations, a more compelling, evolution-based rationale is that breakpoints at the 3’ end of *CIC* preserve functional domain(s) important for activity of the fusion and thus tumorigenesis. In support of this hypothesis, the major exception to C1-domain-retaining breakpoints in CIC::DUX4 fusions is the class of CIC::UTR fusions which are not genuine coding CIC::DUX4 fusions. In these noncanonical fusions, rather than retaining the C1 domain and forming a transactivating fusion with *DUX4*, the breakpoint seems likely to disrupt the C1 domain (and potentially other C-terminal regions) and function as a DUX4-independent, dominant-negative *CIC* mutant as has been hypothesized by others (32). We observed statistically significant but minor activity of such a CIC::UTR fusion when expressed in 293T cells, but further study is warranted, particularly in the context of a potential requirement for haplo-insufficiency and to determine if even mild upregulation of target genes by a potentially dominant-negative CIC::UTR is sufficient for transformation.

We observed through multiple separate cell models that deletion of the C1 domain leads to an attenuated activation of CIC::DUX4 target genes but does not entirely abrogate the CIC::DUX4 transcriptional program. One limitation of interpreting this finding is that not all target genes responded to the same degree, with some maintaining baseline levels following C1-deleted CIC::DUX4 expression while others were significantly upregulated relative to EV conditions. One possible explanation for this depends on recent findings by our group and others that CIC and CIC::DUX4 may additionally bind non-canonical sites on DNA in addition to the preferred octamer (23, 39, 45, 46). If the C1 domain is playing a role in binding DNA in CIC::DUX4, which seems highly likely, then its loss may differentially impact binding to target genes with variable binding motifs in their respective regulatory elements.

A second difficulty in determining whether reduced but significant activation of target genes by C1-deleted CIC::DUX4 is functionally meaningful lies in the field’s lack of understanding of how much CIC::DUX4 expression is ideal for tumorigenesis. This question goes hand-in-hand with the issue of the “Goldilocks Principle” for fusion oncoproteins, as was recently described by Seong and colleagues (40) for EWS::FLI. Our data from CIC::DUX4 stable NIH/3T3 and C2C12 clones seem to converge on the notion that too much full-length CIC::DUX4 is toxic to cells, as the full-length CIC::DUX4 clones expressed much less fusion at the protein level than the weaker C1-deleted CIC::DUX4 clones, despite being driven by an otherwise identical retroviral construct. Indeed, most C2C12 full-length CIC::DUX4 clones did not show detectable fusion protein, suggesting there was selective pressure to silence the transgene. Thus, an improved understanding of endogenous CIC::DUX4 dosage and how it impacts tumor cell survival and death can enhance our understanding of fusion oncoprotein stability and expression.

While exploring phenotypic consequences of deleting the C1 domain, we showed for the first time that CIC::DUX4 can block differentiation of C2C12 cells. This is an exciting finding given the ability of other fusion oncoproteins to cause disease by hijacking differentiation, but we remain unsure if this phenotype is pathogenically relevant. For one, while C1-deleted CIC::DUX4 impaired expression of myogenin, the same transduced cells were unable to initiate tumors *in vivo*. Moreover, while a primitive mesenchymal progenitor is hypothesized to be the cell-of origin for CIC::DUX4 tumors, it is plausible that C2C12 mouse myoblasts are not an accurate cell model to study CIC::DUX4 mediated transformation. Consequently, defining the correct cellular context to study CIC::DUX4 initiation is also an area of importance.

This study has several limitations. First, we did not employ patient-derived cell lines endogenously harboring CIC::DUX4 due to our group’s prior experience attempting to knock out CIC::DUX4 from NCC_CDS1_X1_C1 cells (31) with CRISPR-Cas9, where cell viability was extremely limited upon selection (11), likely due to dependence on the fusion. Consequently, we would expect that mutating the endogenous C1 domain and thus impairing fusion function would lead to loss of cell viability. While the cell lines used here do not directly mimic patient biology, they do reflect proper transcriptional responses to expression of CIC::DUX4 and are easily engineerable, making them useful for mechanistic studies. Second, our *in vivo* model may not completely reflect human biology, since a small number of breakpoints identified in patients are bona fide CIC::DUX4 fusions that do not retain the C1 domain. While the lack of tumorigenesis for C1-deleted CIC::DUX4 expressing C2C12 cells is consistent with their apparent contact inhibited nature from our differentiation experiment, we cannot rule out that C1-deleted CIC::DUX4 may be capable of tumorigenesis in the right cellular context. Third, our breakpoint database is inherently biased by the approaches used to identify breakpoints in patients. Since some CIC-rearranged genes are identified with fixed PCR primers that target only certain parts of the partner genes, this may lead to the unintentional exclusion of breakpoints missed by such approaches.

In summary, we show that the *CIC* C1 domain is necessary for maximal activity of the CIC::DUX4 fusion oncoprotein. This work builds upon pioneering data from wild type CIC and experiments using a fly CIC::DUX4 chimera in *Drosophila* (27) to extend these findings to mammalian and disease relevant settings. Our results demonstrate an example of how paying attention to fusion breakpoints may be informative of which partner gene functional domains are useful to the nascent oncoprotein. Finally, while we do not definitively show that the C1 domain is an essential vulnerability of CIC::DUX4, targeting this key functional domain may be worth further study. Beyond CIC-rearranged tumors, we encourage clinicians and researchers alike to be mindful of reporting fusion oncogene breakpoints in a consistent and accessible manner so that we may use natural evolution of fusions to tell us about their important functional domains.

## Materials and Methods

### Cell Culture

293T (CRL-3216), NIH/3T3 (CRL-1658), and C2C12 (CRL-1772) cells were purchased from ATCC and STR validated using the FTA Human (135-XV) or Mouse (137-XV) Authentication Services (ATCC). All cell lines were grown at 37°C under humidified conditions in 5% CO_2_ incubators in DMEM with high glucose, L-Glutamine, and Sodium Pyruvate (SH30243.02, Cytiva) plus 10% FBS, 100 U/mL Penicillin, and 100 ug/mL Streptomycin. Cell lines were routinely tested for mycoplasma using the e-Myco Plus Mycoplasma PCR Detection Kit (Boca Scientific). Live cells in culture were imaged using a MoticamPro 205A camera mounted on a Trinocular Inverted Microscope (VWR 89404-462).

### CIC-fusion Breakpoint Database and Analysis

Breakpoint coordinates for CIC-rearranged fusion genes described in the literature were manually aligned to reference sequences for *CIC* (NM_015125.5), *DUX4* (NM_001306068.3), *NUTM1* (NM_001284292.2), *LEUTX* (NM_001382345.1), and *FOXO4* (NM_005938.4). When ambiguous bases existed at the breakpoint junction that could not be definitively attributed to either partner, they were assigned to CIC. The full dataset also includes exon-level information for 26 breakpoints where the precise nucleotide coordinates could not be identified, but we advise caution interpreting this data as not all authors use identical reference sequences, and exon numbering is not always consistent across references. While we have done our best to harmonize all data, please refer to the original publications for the most accurate details for specific cases. The full dataset comprised of 108 nucleotide-level and 26 exon-level breakpoints is available for download in Supplemental Dataset S1. Functional domain annotations in Figure 1C and Supplemental Figure S1A were derived from (19, 27, 37, 47). The code used to visualize the data are available at https://github.com/cuylerluck/C1_domain_2024/ (48–51).

### Cloning

HA-CIC::UTR fusion, C1- and HMG-domain deletion and point-mutant constructs were cloned from pcDNA3.1-HA-CIC::DUX4 (a gift from Takuro Nakamura (6)) using the Q5 Site-Directed Mutagenesis Kit (NEB E0554S) with the primers described in Supplemental Dataset S2. The residues deleted for the C1 domain (R1464-M1519, NM_015125.5) and HMG box (I200 – K268, NM_015125.5) deletion mutants were determined from Forés et al., *PLoS Genetics 2017* (27), and Uniprot entry Q96RK0, respectively. The empty vector construct used for 293T experiments was pCMV-Neo-Bam (Addgene 16440), which has the same major functional elements as pcDNA3.1 with the exception of an HSV TK promoter for the Neo resistance gene in place of an SV40 promoter. C1-deleted pMYs-CIC::DUX4-IRES-EGFP was cloned from pMYs-CIC::DUX4-IRES-EGFP (a gift from Takuro Nakamura (52) and Michael Kyba (25)) using the Q5 Site-Directed Mutagenesis Kit (NEB E0554S) with the primers described in Supplemental Dataset S2. The corresponding empty vector pMY-IRES-EGFP was purchased from Addgene (163361). Plasmid sequences were verified via Whole Plasmid Sequencing by Plasmidsaurus/Primordium Labs using Oxford Nanopore Technology with custom analysis and annotation. Select regions were also verified by Sanger sequencing (Quintara Biosciences).

### Plasmid Transfections

For overexpression experiments, typically 1.5 μg of plasmid was reverse transfected into 300,000 293T cells in a well of a 6-well plate using OptiMEM (Gibco) and FuGENE 6 (Promega), at a 2:1 FuGENE:DNA ratio. Transfected cells were typically processed for downstream analysis after approximately 48 hours. For retrovirus generation, 293T cells were forward transfected using the same reagents.

### Western Blot

Adherent cells were washed 3x with DPBS, scraped with RIPA buffer supplemented with Halt protease and phosphatase inhibitors (Thermo Scientific), and incubated on ice for at least 15 minutes. Cell suspensions were then sonicated and centrifuged before supernatant was collected for analysis. Protein samples were normalized, boiled, separated by denaturing electrophoresis on Criterion TGX 4-15% gels (Bio-Rad), transferred to nitrocellulose membranes (Bio-Rad Trans-Blot Turbo), evaluated for transfer by Ponceau S (Sigma-Aldrich) staining, and blocked in 5% BSA in TBS-T for at least one hour. Blots were horizontally cut to use for separate targets. Primary antibodies were diluted in blocking buffer and incubated overnight followed by washing with TBS-T, and secondary antibodies were similarly diluted and incubated for one hour. After washing with TBS-T, imaging was performed on a Bio-Rad ChemiDoc Touch using ECL Prime reagent (Millipore Sigma) by briefly drying the blot, submerging the blot in ECL Prime mixture, dabbing excess solution off, and imaging. When required, brightness/contrast was adjusted either on the ChemiDoc or using Image Lab (Bio-Rad). Antibodies used and their dilutions were: HSP90 (CST 4874S, 1:1000), HA-tag clone C29F4 (CST 3724S, 1:1000), ETV5 (CST 16274S, 1:1000), DUX4 clone P4H2 (Invitrogen MA5-16147, 1:1000-1:2000), GFP (CST 2956S, 1:1000-1:5000), Myogenin clone F5D (Invitrogen 14-5643-82, 1:2000), HRP-linked Rabbit IgG (CST 7074, 1:3000), and HRP-linked Mouse IgG (CST 7076, 1:3000).

### RT-qPCR

RNA was extracted from adherent cells with the RNeasy Mini kit (Qiagen) per manufacturer’s protocol. 1 ug RNA per sample was used as input to the SensiFAST cDNA Synthesis Kit (Bioline). RT-qPCR was performed in technical quadruplicate with TaqMan Fast Advanced Master Mix (Applied Biosystems) and TaqMan probes on an Applied Biosystems StepOnePlus. Probes used were *GAPDH* (Hs02758991_g1), *ETV1* (Hs00951951_m1), *ETV4* (Hs00383361_g1), *ETV5* (Hs00927557_m1), and *DUSP6* (Hs04329643_s1). Gene expression was normalized to GAPDH using the 2^(Ct^ ^housekeeping−Ct^ ^experimental^ ^gene)^ method. To combine replicates for statistical analysis, replicates were first individually normalized with 0% defined as the average of the EV condition, and 100% defined as the average of the HA-CIC::DUX4 condition. Then, technical quadruplicates were averaged for each replicate and aggregated. Statistical analysis was performed using one-way ANOVA and Šidák’s multiple comparisons test. GraphPad Prism (version 10.2.3) was used for statistical analysis.

### Retroviral Transductions and Clonal Cell Line Generation

Ecotropic retrovirus was generated in 293T cells by cotransfection of one of the transfer vectors (pMY-IRES-EGFP, pMYs-CIC::DUX4-IRES-EGFP, or pMYs-CIC::DUX4-IRES-EGFP dC1) and the packaging vector pCL-Eco (Novus Biologicals NBP2-29540). NIH/3T3 or C2C12 cells were infected by addition of viral supernatant with 10 ug/mL Polybrene (Millipore Sigma) directly on top of cells and were bulk sorted for EGFP+ cells 48 hours after infection with a FACSAria II (BD Biosciences). To generate clonal cell lines, a FACSAria III (BD Biosciences) was used to further sort single EGFP+ cells into individual wells of a 96-well plate. Wells with a single colony confirmed by microscopic inspection were expanded for freezing and evaluated for transgene expression by western blot before further use.

### Luciferase Reporter Assay

In a 96-well plate, 10,000 293T cells per well were reverse transfected as described above with 50 ng each of experimental plasmid (EV or HA-CIC::DUX4/mutants) and a GoClone *ETV5* promoter-RenSP construct (SwitchGear Genomics S722373) containing the consensus CIC binding sequence TGAATGAA. Transfections were performed in technical quadruplicate. After approximately 48 hours, cells were processed with the Lightswitch Assay Kit (Active Motif LS010) and luciferase signal was read for 1.5 seconds on a SpectraMax M5 (Molecular Devices). To combine replicates for statistical analysis, replicates were first individually normalized with 0% defined as the average of the EV condition, and 100% defined as the average of the HA-CIC::DUX4 condition. Then, technical quadruplicates were averaged for each replicate and aggregated. Statistical analysis was performed using one-way ANOVA and Šidák’s multiple comparisons test. GraphPad Prism (version 10.2.3) was used for statistical analysis.

### Immunofluorescence

For transfected 293T, cells were first reverse transfected as described above with appropriate plasmids. The next day, transfected cells were passaged onto collagen I-coated (50ug/ml, Corning 354236) 18mm coverslips. Approximately 48 hours after transfection, cells were fixed with 4% PFA in PBS for 10 minutes at 37C, briefly washed once with DPBS, quenched with 100 mM glycine for 30 minutes at room temperature, briefly washed twice with DPBS, permeabilized with 0.2% Triton X-100 for 15 minutes at room temperature, washed three times with DPBS, and blocked in 2% BSA overnight at 4°C. Coverslips were incubated with primary antibody (HA-tag clone C29F4, CST 3724S, dilution 1:300) for two hours at room temperature, washed three times with DPBS, and again incubated with a secondary mixture of 1:10,000 rhodamine phalloidin (Invitrogen R415), 1:1000 DAPI (Thermo Fisher D1306), and 1:300 anti-Rabbit IgG Alexa 647 (Invitrogen A-21245) for one hour at room temperature. Coverslips were washed three times with DPBS and mounted in ProLong Glass (Invitrogen P36984). Cells were imaged on a Yokogawa CSU-X1 spinning disk confocal on a Nikon Ti-E microscope with a 60x/1.45NA lens (Nikon) using NIS Elements AR (v5.21.03, Nikon) software at consistent exposure times and laser powers. Images were processed with FIJI/ImageJ (https://github.com/fiji/fiji). Briefly, all confocal planes were converted to single images by maximum intensity projection and brightness/contrast was consistently adjusted between all images.

For transduced clonal NIH/3T3, cells were seeded onto poly-L-lysine (0.01%, Sigma-Aldrich P4707-50ML) coated coverslips and allowed to adhere overnight. Cells were then similarly prepared as above but blocked for one hour at room temperature, prepared without a primary antibody step, and used 1:400 rhodamine phalloidin (Invitrogen R415) and 1:2000 DAPI (Thermo Fisher Scientific D1306). Cells were imaged on a Zeiss Axio Imager 2 with Zeiss ZEN 2 (blue edition, version 2.0.0.0) software and a 20x/0.8 Plan-APOCHROMAT air objective (Zeiss) at consistent exposure times and laser powers, with direct detection of EGFP fluorescence. Images were processed as described above, without the maximum intensity projection step.

### RNA Sequencing and Analysis

150,000 cells per cell line were seeded in wells of a 6-well plate and allowed to grow for 48 hours. After 48 hours, RNA was extracted using the RNeasy Mini kit (Qiagen) including an on-column DNase digest (Qiagen) and an additional short Buffer RPE wash. RNA samples were additionally processed with the Monarch RNA Cleanup Kit (NEB), analyzed for integrity with a TapeStation (Agilent), and submitted to Novogene for library preparation and sequencing. Samples were polyA-enriched and directional libraries were prepared using the NEBNext Ultra II Directional Library Prep Kit for Illumina (NEB). Libraries were PE150 sequenced on a NovaSeq 6000 (Illumina) with approximately 10G of data per sample.

FASTQ files showed no concerning quality indications per both the Novogene report and our own analysis with FastQC (53). Full code describing processing and analysis of the data is available at https://github.com/cuylerluck/C1_domain_2024/. Briefly, FASTQ files were aligned to the mouse GRCm39 genome with STAR (54), including quantitation of gene counts using the option -- quantMode GeneCounts. Uniquely mapped read rates were between 92% and 94% for all samples. The grep function was used to manually verify the presence of proper transgenes in each sample. A custom R script (version 4.2.2) (55–62), also available at https://github.com/cuylerluck/C1_domain_2024/, was used to perform differential gene expression analysis. Briefly, column 4 from STAR GeneCount output was merged between all samples. Then, edgeR (version 3.40.2) (63–66) was used for differential expression analysis with a GLM method and quasi-likelihood F-tests, with resulting p-values then FDR-corrected. log2(counts per million) values were extracted from the TMM-normalized dataset using the cpm function. The differential expression output for both the CD4 vs. EV and dC1 vs EV comparisons are available in Supplemental Datasets S3 and S4. The raw FASTQ files, log2(cpm) data, and counts data have been deposited in NCBI’s Gene Expression Omnibus (67) and are accessible through GEO Series accession number GSE269295 (https://www.ncbi.nlm.nih.gov/geo/query/acc.cgi?acc=GSE269295).

### C2C12 Differentiation Assay

Clonal C2C12 cell lines were seeded at 175,000 cells per well in wells of identically prepared 6-well plates, one plate per time point. The next day, media was removed, gently rinsed once with DPBS, and differentiation media was added (growth DMEM as defined above, but with 2% horse serum [Thermo Fisher Scientific 16050130] in place of FBS). The day 0 plate was immediately imaged and harvested for protein as described above. Differentiation media was changed 2 days after differentiation began, and the day 4 plate was imaged. The day 4 plate was imaged and harvested another 2 days later, while differentiation media was replaced on the day 6 plate. Finally, the day 6 plate was imaged and harvested after another 2 days. Protein samples were analyzed by western blot as described above.

For the siRNA + differentiation experiment, 175,000 clonal C2C12 cells per well were plated in 6-well plates. The next day, one plate was briefly rinsed once with DPBS, switched to differentiation media, and immediately harvested for protein. The other plate was briefly rinsed once with DPBS, switched to differentiation media, and cells were forward transfected with 100 pmol of appropriate siRNA Dharmacon ON-TARGETplus SMARTPools, either non-targeting (Horizon D-001810-10-05), or human siCIC (Horizon L-015185-01-0020) using Lipofectamine RNAiMAX (Invitrogen). Differentiation media was then changed two and four days after transfection, and cells were harvested for protein five days after infection. Protein samples were analyzed by western blot as described above. The sequences for the siRNA pools were as follows. Non-targeting: UGGUUUACAUGUCGACUAA, UGGUUUACAUGUUGUGUGA, UGGUUUACAUGUUUUCUGA, UGGUUUACAUGUUUUCCUA. Human siCIC: GCUUAGUGUAUUCGGACAA, CGGCGCAAGAGACCCGAAA, GAGAAGCCGCAAUGAGCGA, CGAGUGAUGAGGAGCGCAU. None of the human siCIC sequences are perfect matches for the mouse short *CIC* isoform (NM_027882.4), each having one to three mismatches with mouse *CIC*.

### Hanging Drop Assay

Cells were diluted to 1.5e6 viable cells per mL and 5 uL droplets (7,500 cells per droplet) were placed on the underside of the lid of a 10cm tissue culture dish filled with 10 mL of sterile DPBS. 12 droplets were placed per plate. Droplets were incubated for 24 hours, after which the lid was inverted droplet side-up and imaged using a MoticamPro 205A camera mounted on a Trinocular Inverted Microscope (VWR 89404-462).

### *In Vivo* Tumorigenesis

Five to six week old female nude (NU/J) mice were purchased from Jackson Laboratory (RRID:IMSR_JAX:002019). Specific pathogen-free conditions and facilities were approved by the American Association for Accreditation of Laboratory Animal Care. Mouse experiments followed the reviewed and approved UCSF IACUC protocol AN194620-01C. For subcutaneous flank injection, cell lines were prepared to 8 million cells per 500 uL DPBS and 25 uL of cell suspension was mixed 1:1 with Matrigel (Corning 356234) to achieve 400,000 cells per 50 uL injection. For each of the three cell lines tested, five mice per group were anesthetized with isoflurane and subcutaneously injected in both flanks. Mice were monitored for tumor size three times a week starting four days post-injection, with measurable lesions evaluated with a digital caliper. Tumor volume was calculated by the (pi/6)*length*width*height method. The R script and raw data used to produce the plot of tumor size versus time is available at https://github.com/cuylerluck/C1_domain_2024/. Upon sacrifice, collectable tumor tissue was resected from euthanized mice and stored on ice, quickly frozen on dry ice upon return to the lab, and stored at −80C.

Protein lysates were collected from tumor tissue by mortar and pestle in RIPA buffer supplemented with protease and phosphatase inhibitors, followed by sonication and centrifugation as above. Cell line control lysate was prepared from cells plated for 48 hours as described above. Western blotting was performed as described above.

For RNA extraction, tumor tissue was homogenized using a Precellys Evolution (Bertin Technologies) with High Impact Zirconium beads (Benchmark Scientific D1032-30). RNA was then extracted using the RNeasy Mini kit (Qiagen) including an on-column DNase digest (Qiagen) and 500 ng RNA per sample was used as input to the SensiFAST cDNA Synthesis Kit (Bioline), including -RT controls. Cell line control cDNA was similarly prepared from cells plated for 48 hours. PCR was performed on the cDNA samples using Phusion polymerase (NEB). EGFP reactions used HF buffer, annealing temperature 66C, 35 cycles, and primers EGFP_FWD (5’ taaacggccacaagttcagcg 3’) and EGFP_REV (5’ cagctcgatgcggttcaccag 3’). CIC::DUX4 reactions used GC buffer, annealing temperature 68C, 40 cycles, and primers CIC_FWD (5’ agaagccgaggacgtgcttgg 3’) and DUX4_REV (5’ gttgcgcctgctgcagaaac 3’). PCR products were electrophoresed on a 2% agarose gel made with Ethidium Bromide (Fisher Scientific) and imaged on a ChemiDoc Touch (Bio-Rad).

## Acknowledgments

C.L. acknowledges funding from the UCSF BMS Graduate Program Training Grant T32GM136547-01 (NIGMS), and the UCSF Discovery Fellows Program. K.A.J. acknowledges funding from Tobacco-Related Disease Research Program Predoctoral Fellowship T33DT6442. This material is based upon work supported by the National Science Foundation Graduate Research Fellowship Program under Grant No. 2038436 (to K.A.J.). Any opinions, findings, and conclusions or recommendations expressed in this material are those of the author(s) and do not necessarily reflect the views of the National Science Foundation. R.A.O. acknowledges funding from an NCI grant (R37 CA255453), the Children’s Cancer Research Fund (CCRF), and Cookies for Kids’ Cancer. We acknowledge the PFCC (RRID:*SCR_018206*) supported in part by Grant NIH P30 DK063720 and by the NIH S10 Instrumentation Grant S10 1S10OD021822-01 for their assistance and expertise with flow cytometry. We thank Takuro Nakamura and Michael Kyba for sharing of the original pcDNA3.1-HA-CIC::DUX4 and pMYs-CIC::DUX4-IRES-EGFP constructs. pMY-IRES-eGFP was a gift from Louis Ates & Teunis Geijtenbeek (Addgene plasmid #163361; http://n2t.net/addgene:163361; RRID:Addgene_163361) (68). pCMV-Neo-Bam was a gift from Bert Vogelstein (Addgene plasmid #16440; http://n2t.net/addgene:16440; RRID:Addgene_16440) (69). Many thanks to Kevin Shannon, Alejandro Sweet-Cordero, and Dave Toczyski for their valuable input as this project developed.

## Notes

https://www.ncbi.nlm.nih.gov/geo/query/acc.cgi?acc=GSE269295

https://github.com/cuylerluck/C1_domain_2024/

### Competing Interest Statement

The authors have declared no competing interest.

## References

1. S. Zhang, et al., PAX3-FOXO1 coordinates enhancer architecture, eRNA transcription, and RNA polymerase pause release at select gene targets. Mol. Cell 82, 4428–4442.e7 (2022).

2. B. E. Gryder, et al., PAX3-FOXO1 establishes myogenic super enhancers and confers BET bromodomain vulnerability. Cancer Discov. 7, 884–899 (2017).

3. N. Riggi, et al., EWS-FLI1 Utilizes Divergent Chromatin Remodeling Mechanisms to Directly Activate or Repress Enhancer Elements in Ewing Sarcoma. Cancer Cell 26, 668– 681 (2014).

4. G. Boulay, et al., Cancer-Specific Retargeting of BAF Complexes by a Prion-like Domain. Cell 171, 163–178.e19 (2017).

5. M. J. McBride, et al., The SS18-SSX Fusion Oncoprotein Hijacks BAF Complex Targeting and Function to Drive Synovial Sarcoma. Cancer Cell 33, 1128–1141.e7 (2018).

6. M. Kawamura-Saito, et al., Fusion between CIC and DUX4 up-regulates PEA3 family genes in Ewing-like sarcomas with t(4;19)(q35;q13) translocation. Hum. Mol. Genet. 15, 2125–2137 (2006).

7. A. S. Brohl, et al., The Genomic Landscape of the Ewing Sarcoma Family of Tumors Reveals Recurrent STAG2 Mutation. PLoS Genet. 10 (2014).

8. J. F. Shern, et al., Comprehensive genomic analysis of rhabdomyosarcoma reveals a landscape of alterations affecting a common genetic axis in fusion-positive and fusion-negative tumors. Cancer Discov. 4, 216–231 (2014).

9. Y. Asante, et al., PAX3-FOXO1 uses its activation domain to recruit CBP/P300 and shape RNA Pol2 cluster distribution. Nat. Commun. 14 (2023).

10. P. Y. P. Lam, J. E. Sublett, A. D. Hollenbach, M. Roussel, The Oncogenic Potential of the Pax3-FKHR Fusion Protein Requires the Pax3 Homeodomain Recognition Helix but Not the Pax3 Paired-Box DNA Binding Domain. Mol. Cell. Biol. 19, 7287–7287 (1999).

11. Y. K. Lin, W. Wu, R. K. Ponce, J. W. Kim, R. A. Okimoto, Negative MAPK-ERK regulation sustains CIC-DUX4 oncoprotein expression in undifferentiated sarcoma. Proc. Natl. Acad. Sci. (2020). 10.1073/pnas.2009137117.

12. R. K. M. Ponce, C. Luck, R. A. Okimoto, Molecular and therapeutic advancements in Capicua (CIC)-rearranged sarcoma. Front. Cell Dev. Biol. 12, 1–11 (2024).

13. S. Watson, et al., Transcriptomic definition of molecular subgroups of small round cell sarcomas. J. Pathol. 245, 29–40 (2018).

14. M. Berger, et al., Genomic EWS-FLI1 Fusion Sequences in Ewing Sarcoma Resemble Breakpoint Characteristics of Immature Lymphoid Malignancies. PLoS One 8, 1–7 (2013).

15. C. Vicente-García, et al., Regulatory landscape fusion in rhabdomyosarcoma through interactions between the PAX3 promoter and FOXO1 regulatory elements. Genome Biol. 18 (2017).

16. G. Jiménez, A. Guichet, A. Ephrussi, J. Casanova, Relief of gene repression by Torso RTK signaling: Role of capicua in Drosophila terminal and dorsoventral patterning. Genes Dev. 14, 224–231 (2000).

17. S. T. Ahmad, et al., Capicua regulates neural stem cell proliferation and lineage specification through control of Ets factors. Nat. Commun. 10, 1–17 (2019).

18. Y. Ren, et al., CIC is a mediator of the ERK1/2-DUSP6 negative feedback loop. iScience 105398 (2020). 10.1016/j.isci.2020.101635.

19. K. Dissanayake, et al., ERK/p90RSK/14-3-3 signalling has an impact on expression of PEA3 Ets transcription factors via the transcriptional repressor capicúa. Biochem. J. 433, 515–525 (2011).

20. V. G. LeBlanc, et al., Comparative transcriptome analysis of isogenic cell line models and primary cancers links capicua (CIC) loss to activation of the MAPK signalling cascade. J. Pathol. 242, 206–220 (2017).

21. S. Weissmann, et al., The tumor suppressor CIC directly regulates MAPK pathway genes via histone deacetylation. Cancer Res. 78, 4114–4125 (2018).

22. J. W. Kim, R. K. Ponce, R. A. Okimoto, Capicua in Human Cancer. Trends in Cancer 1–10 (2020). 10.1016/j.trecan.2020.08.010.

23. P. G. Hendrickson, et al., Spontaneous expression of the CIC::DUX4 fusion oncoprotein from a conditional allele potently drives sarcoma formation in genetically engineered mice. Oncogene (2024). 10.1038/s41388-024-02984-8.

24. R. A. Okimoto, et al., CIC-DUX4 oncoprotein drives sarcoma metastasis and tumorigenesis via distinct regulatory programs. J. Clin. Invest. 129, 3401–3406 (2019).

25. D. Bosnakovski, et al., Inactivation of the CIC-DUX4 oncogene through P300/CBP inhibition, a therapeutic approach for CIC-DUX4 sarcoma. Oncogenesis 10, 1–11 (2021).

26. C. R. Antonescu, et al., Sarcomas With CIC-rearrangements Are a Distinct Pathologic Entity With Aggressive Outcome A Clinicopathologic and Molecular Study of 115 Cases. Am J Surg Pathol 41, 941–949 (2017).

27. M. Forés, et al., A new mode of DNA binding distinguishes Capicua from other HMG-box factors and explains its mutation patterns in cancer. PLoS Genet. 13, 1–22 (2017).

28. J. Webb, et al., Molecular basis of DNA recognition by the HMG-box-C1 module of Capicua Running title: DNA binding of CIC HMG-box and C1 domains. bioRxiv (2022). 10.1101/2022.03.28.485992.

29. S. Nakai, et al., Establishment of a novel human CIC-DUX4 sarcoma cell line, Kitra-SRS, with autocrine IGF-1R activation and metastatic potential to the lungs. Sci. Rep. 9, 1–6 (2019).

30. Y. Yoshimatsu, et al., Establishment and characterization of NCC-CDS2-C1: a novel patient-derived cell line of CIC-DUX4 sarcoma. Hum. Cell 33, 427–436 (2020).

31. R. Oyama, et al., Generation of novel patient-derived CIC-DUX4 sarcoma xenografts and cell lines. Sci. Rep. 7, 1–13 (2017).

32. I. Panagopoulos, et al., Chromosome Translocation t(10;19)(q26;q13) in a CIC-sarcoma. In Vivo (Brooklyn). 37, 57–69 (2023).

33. A. Yoshida, et al., CIC break-apart fluorescence in-situ hybridization misses a subset of CIC–DUX4 sarcomas: a clinicopathological and molecular study. Histopathology 71, 461– 469 (2017).

34. M. Gambarotti, et al., CIC–DUX4 fusion-positive round-cell sarcomas of soft tissue and bone: a single-institution morphological and molecular analysis of seven cases. Histopathology 69, 624–634 (2016).

35. S. Cocchi, et al., CIC rearranged sarcomas: A single institution experience of the potential pitfalls in interpreting CIC FISH results. Pathol. Res. Pract. 231 (2022).

36. Y. Tsukamoto, et al., Primary undifferentiated small round cell sarcoma of the deep abdominal wall with a novel variant of t(10;19) CIC-DUX4 gene fusion. Pathol. Res. Pract. 213, 1315–1321 (2017).

37. H. Mitsuhashi, et al., Functional domains of the FSHD-associated DUX4 protein. Biol. Open 7 (2018).

38. S. H. Choi, et al., DUX4 recruits p300/CBP through its C-terminus and induces global H3K27 acetylation changes. Nucleic Acids Res. 44, 5161–5173 (2016).

39. A. Papagianni, et al., Capicua controls Toll/IL-1 signaling targets independently of RTK regulation. Proc. Natl. Acad. Sci. U. S. A. 115, 1807–1812 (2018).

40. B. K. A. Seong, et al., TRIM8 modulates the EWS/FLI oncoprotein to promote survival in Ewing sarcoma. Cancer Cell 39, 1262–1278.e7 (2021).

41. V. Andrés, K. Walsh, Myogenin expression, cell cycle withdrawal, and phenotypic differentiation are temporally separable events that precede cell fusion upon myogenesis. J. Cell Biol. 132, 657–666 (1996).

42. A. Wernig, et al., Formation of new muscle fibres and tumours after injection of cultured myogenic cells. J. Neurocytol. 20, 982–997 (1991).

43. Y. Hamamori, B. Samal, J. Tian, L. Kedes, Persistent Erythropoiesis by Myoblast Transfer of Erythropoietin cDNA. Hum. Gene Ther. 5, 1349–1356 (1994).

44. J. E. Morgan, S. E. Moore, F. S. Walsh, T. A. Partridge, Formation of skeletal muscle in vivo from the mouse C2 cell line. J. Cell Sci. 102, 779–787 (1992).

45. J. W. Kim, et al., Capicua suppresses YAP1 to limit tumorigenesis and maintain drug sensitivity in human cancer. Cell Rep. 41, 111443 (2022).

46. N. J. Thomas, et al., Mapping chromatin state and transcriptional response in CIC-DUX4 undifferentiated round cell sarcoma. bioRxiv Prepr. Serv. Biol. 1–36 (2023). 10.1101/2023.10.11.561932.

47. A. S. Futran, S. Kyin, S. Y. Shvartsman, A. J. Link, Mapping the binding interface of ERK and transcriptional repressor Capicua using photocrosslinking. PNAS 112 (2015).

48. Z. Gu, R. Eils, M. Schlesner, Complex heatmaps reveal patterns and correlations in multidimensional genomic data. Bioinformatics 32, 2847–2849 (2016).

49. Z. Gu, L. Gu, R. Eils, M. Schlesner, B. Brors, circlize implements and enhances circular visualization in R. Bioinformatics 30, 2811–2812 (2014).

50. E. Neuwirth, RColorBrewer: ColorBrewer Palettes. (2022). Available at: https://cran.r-project.org/package=RColorBrewer.

51. D. Attali, C. Baker, ggExtra: Add Marginal Histograms to “ggplot2”, and More “ggplot2” Enhancements. (2022). Available at: https://cran.r-project.org/package=ggExtra.

52. T. Yoshimoto, et al., CIC-DUX4 induces small round cell sarcomas distinct from ewing sarcoma. Cancer Res. 77, 2927–2937 (2017).

53. S. Andrews, FastQC. (2010). Available at: https://www.bioinformatics.babraham.ac.uk/projects/fastqc/.

54. A. Dobin, et al., STAR: Ultrafast universal RNA-seq aligner. Bioinformatics 29, 15–21 (2013).

55. R Core Team, R: A Language and Environment for Statistical Computing. (2022). Available at: https://www.r-project.org/.

56. M. Dowle, A. Srinivasan, data.table: Extension of ‘data.framè. (2023). Available at: https://cran.r-project.org/package=data.table.

57. H. Wickham, D. Vaughan, M. Girlich, tidyr: Tidy Messy Data. (2023). Available at: https://cran.r-project.org/package=tidyr.

58. H. Wickham, R. François, L. Henry, K. Müller, D. Vaughan, dplyr: A Grammar of Data Manipulation. (2023). Available at: https://cran.r-project.org/package=dplyr.

59. H. Wickham, ggplot2: Elegant Graphics for Data Analysis. (2016). Available at: https://ggplot2.tidyverse.org.

60. R. Kolde, pheatmap: Pretty Heatmaps. (2019). Available at: https://cran.r-project.org/package=pheatmap.

61. K. Slowikowski, ggrepel: Automatically Position Non-Overlapping Text Labels with “ggplot2.” (2023). Available at: https://cran.r-project.org/package=ggrepel.

62. T. L. Pedersen, patchwork: The Composer of Plots. (2022). Available at: https://cran.r-project.org/package=patchwork.

63. M. D. Robinson, D. J. McCarthy, G. K. Smyth, edgeR : a Bioconductor package for differential expression analysis of digital gene expression data. Bioinformatics 26, 139– 140 (2010).

64. M. D. Robinson, A. Oshlack, A scaling normalization method for differential expression analysis of RNA-seq data. Genome Biol. 11, R25 (2010).

65. D. J. McCarthy, Y. Chen, G. K. Smyth, Differential expression analysis of multifactor RNA-Seq experiments with respect to biological variation. Nucleic Acids Res. 40, 4288–4297 (2012).

66. A. T. L. Lun, Y. Chen, G. K. Smyth, It’s DE-licious: A Recipe for Differential Expression Analyses of RNA-seq Experiments Using Quasi-Likelihood Methods in edgeR. Methods Mol. Biol. 1418, 391–416 (2016).

67. R. Edgar, M. Domrachev, A. E. Lash, Gene Expression Omnibus: NCBI gene expression and hybridization array data repository. Nucleic Acids Res. 30, 207–210 (2002).

68. L. E. H. Van der Donk, et al., An optimized retroviral toolbox for overexpression and genetic perturbation of primary lymphocytes. Biol. Open 11, 1–11 (2022).

69. S. J. Baker, S. Markowitz, E. R. Fearon, J. K. V Willson, B. Vogelstein, Suppression of Human Colorectal Carcinoma Cell Growth by Wild-Type p53. Science (80-.). 249, 912– 915 (1990).

