## Supplemental Figures S1-S6 for "The *Capicua* C1 Domain is Required for Full Activity of the CIC::DUX4 Fusion Oncoprotein"

The supplemental data contains six figures and four datasets.

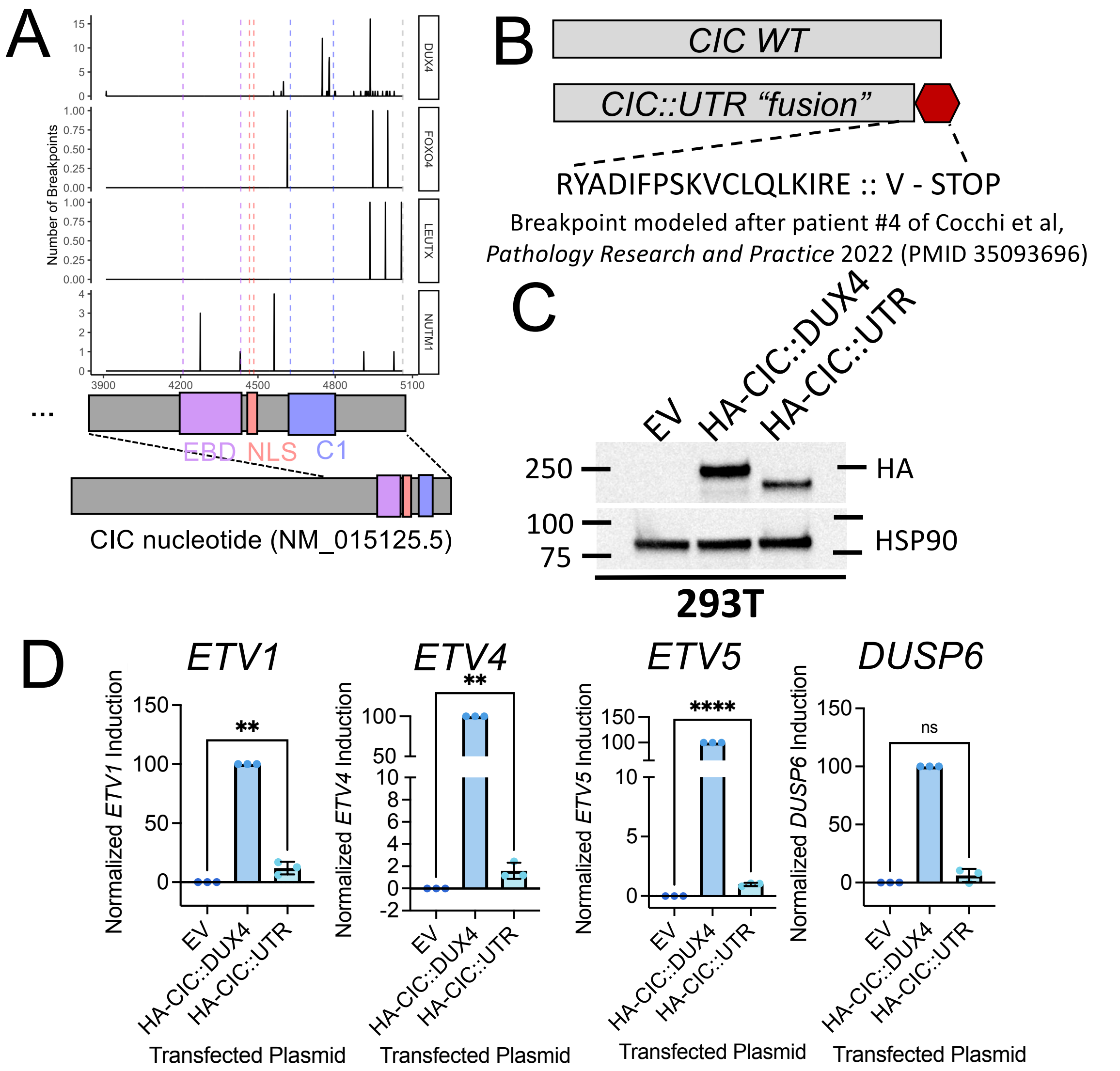


**Supplemental Figure S1.** *CIC* breakpoints are mildly variable across different 3' partner genes, and piloting of a *CIC*::UTR "fusion" model. (A) Histogram of *CIC* breakpoints across four 3' partner genes for a total of 90 breakpoints using RNA data where both partner breakpoints were available. Breakpoint numbers by partner gene: DUX4 = 74, NUTM1 = 10, LEUTX = 3, FOXO4 = 3. EBD = ERK-binding domain, NLS = nuclear localization signal, C1 = C1 domain. (B) Schematic for design of CIC::UTR construct, WT CIC is shown for visual comparison but HA-CIC::UTR was cloned from an HA-CIC::DUX4 plasmid. (C) Immunoblot of 293T cells approximately 48 hours after transfection with empty vector (EV) or the labeled constructs, data representative of three independent experiments. (D) Normalized RT-qPCR measurement of target gene induction in 293T cells approximately 48 hours after transfection with EV or the labeled constructs. Each data point represents the mean of one of three independent experiments, error bars indicate standard deviation, **** = p < 0.0001, and ** = p < 0.01 by one-way ANOVA and Šidák's multiple comparisons test.

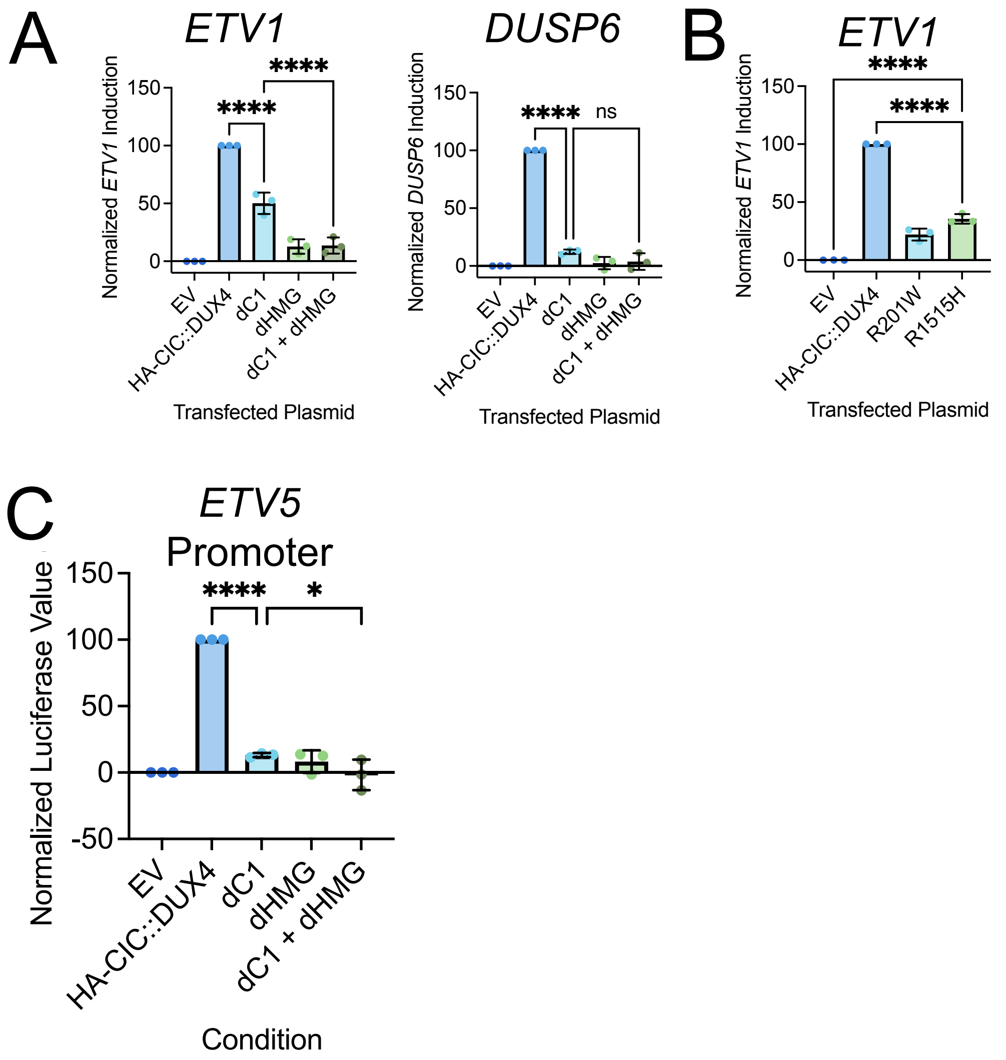


**Supplemental Figure S2.** Deletion or point mutation of the C1 domain result in attenuated target gene activation and ETV5-promoter engagement by CIC::DUX4. (A and B) Normalized RT-qPCR measurement of target gene induction in 293T cells approximately 48 hours after transfection with EV or the labeled constructs. Each data point represents the mean of one of three independent experiments, error bars indicate standard deviation, **** = $p < 0.0001$, and ns = $p > 0.05$ by one-way ANOVA and Šidák's multiple comparisons test. (C) Normalized luciferase measurement of target gene induction in 293T cells approximately 48 hours after cotransfection with EV or the labeled constructs along with an ETV5-promoter-luciferase reporter. Each data point represents the mean of one of three independent experiments, error bars indicate standard deviation, **** = $p < 0.0001$, and * = $p \leq 0.05$ by one-way ANOVA and Šidák's multiple comparisons test.

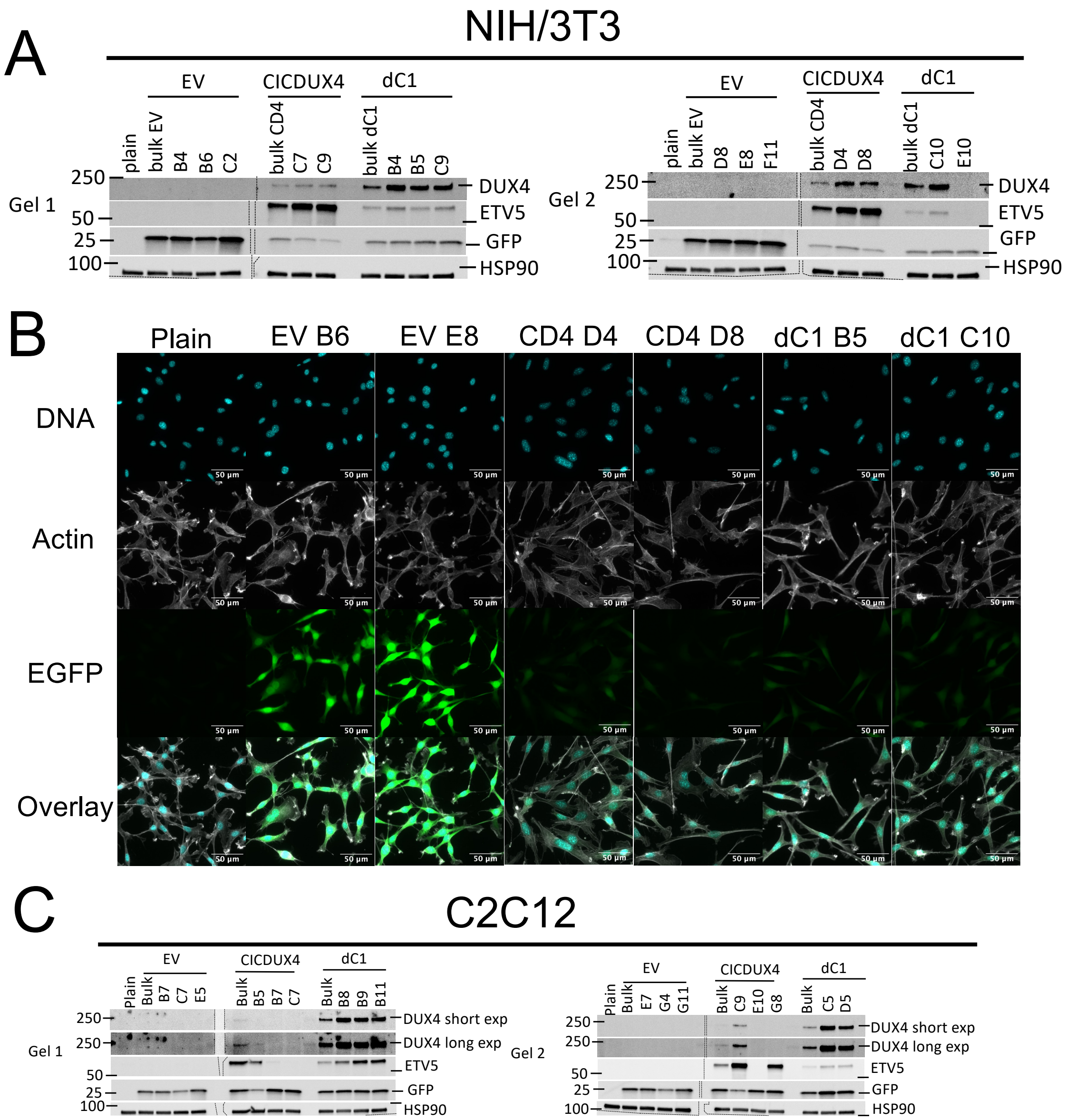


**Supplemental Figure S3.** Screening and validation of transduced NIH/3T3 and C2C12 clonal cell lines. (A) Immunoblot of untransduced (plain), polyclonal (bulk), or clonal (number-letter) NIH/3T3 cells. Abbreviated labels for transgenes use the same naming scheme as those in Figure 3A. Dashed lines indicate where blots were physically cut due to space limitations. Blank lanes had ladder run in them. This immunoblot for clone screening was performed once. (B) Epifluorescence microscopy of untransduced (plain) or selected clonal NIH/3T3 cells. DNA visualized with DAPI, actin visualized with rhodamine-phalloidin, 20x objective used for imaging, scale bars indicate 50 μm, representative cells chosen from one experiment. (C) Immunoblot of untransduced (plain), polyclonal (bulk), or clonal (number-letter) C2C12 cells. Abbreviated labels for transgenes use the same naming scheme as those in Figure 3A. “short exp” or “long exp” indicates different exposure times. Dashed lines indicate where blots were physically cut due to space limitations. Blank lanes had ladder run in them. This immunoblot for clone screening was performed once.

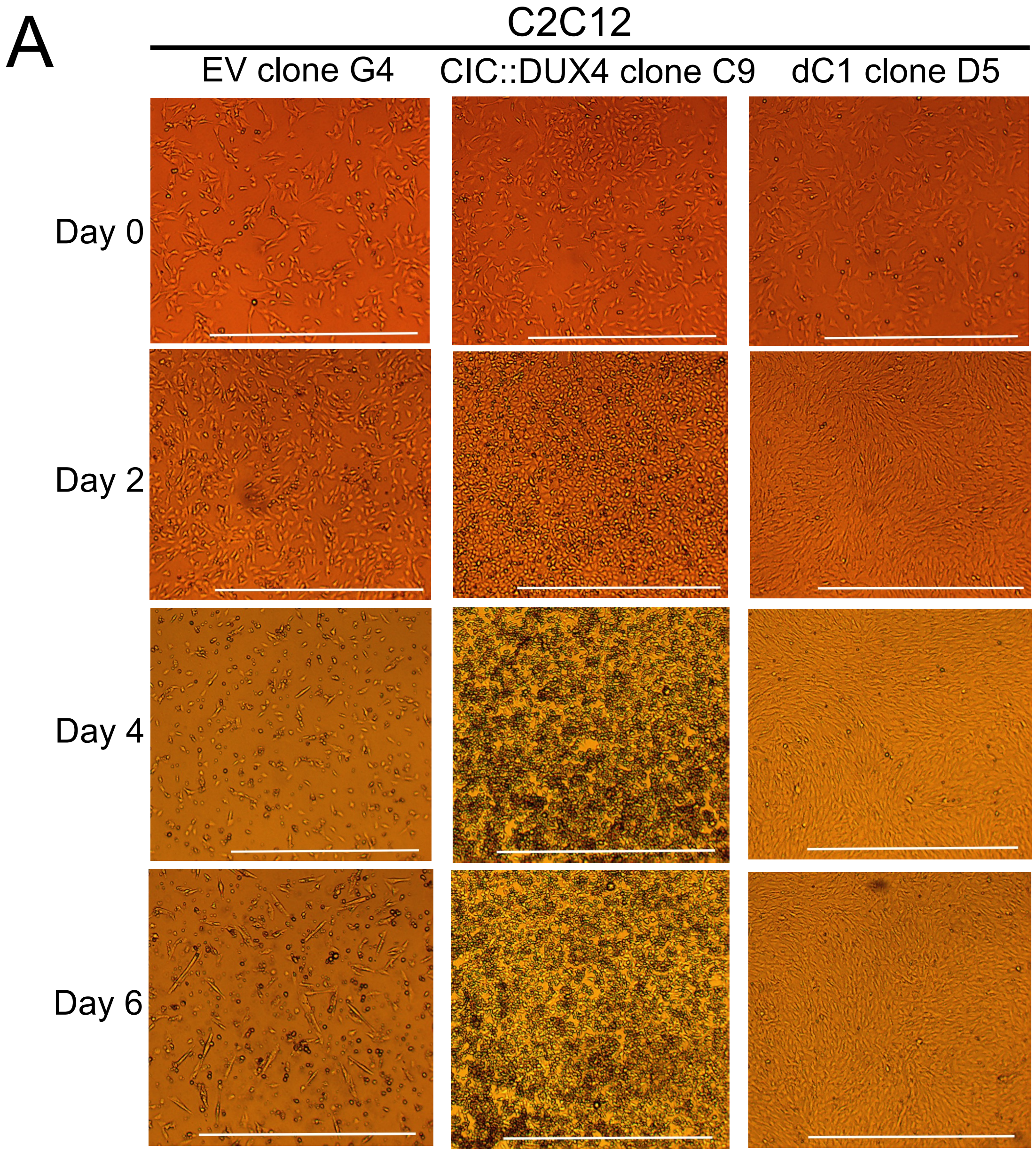


**Supplemental Figure S4.** C1-intact or deleted CIC::DUX4 expression alters growth patterns of clonal C2C12 cells. (A) Clonal C2C12 cell lines expressing the indicated transduced constructs imaged at the indicated timepoints during a differentiation experiment. Images are from different regions of wells and/or different identically prepared plates between timepoints. Imaged with a 4x objective, scale bar represents 1mm. Representative of two independent experiments.

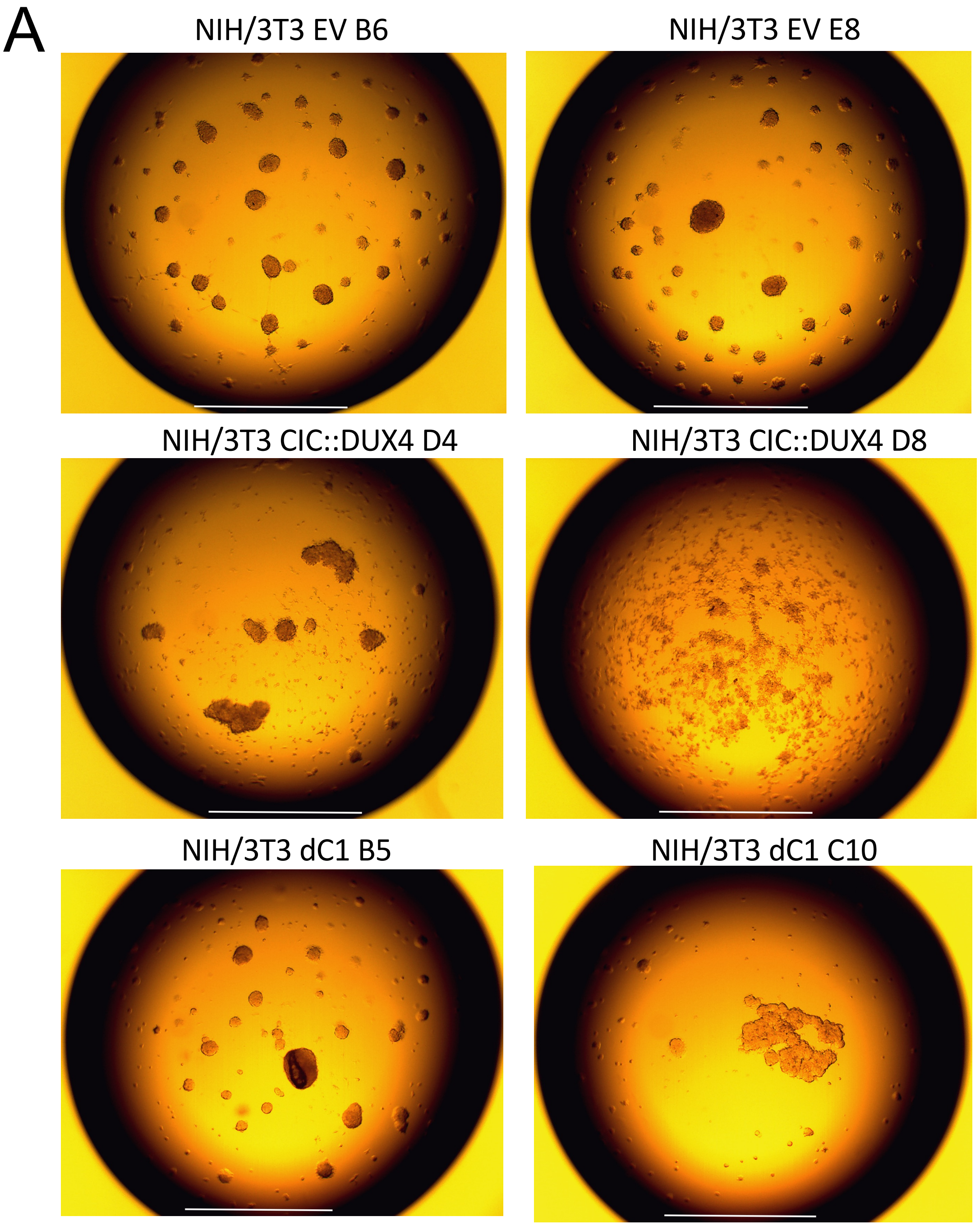


**Supplemental Figure S5.** Full-length or C1-deleted CIC::DUX4 expression alters 3D growth in NIH/3T3 clones. (A) Images of the indicated NIH/3T3 clones after approximately 24 hours in a hanging drop assay. Imaged with a 4x objective, scale bar represents 1mm. Representative drops are shown from three independent experiments, each with 12 drops per condition.

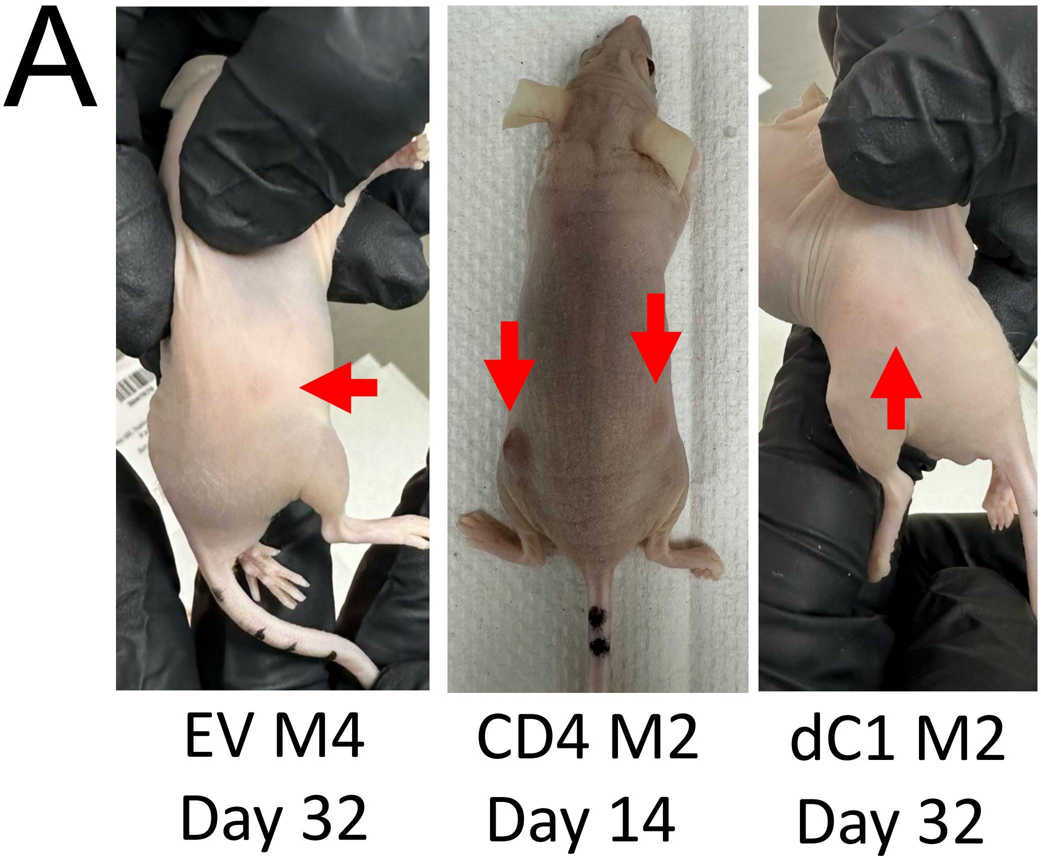


**Supplemental Figure S6.** Only full-length CIC::DUX4 expressing C2C12 cells are capable of forming overt tumors in a nude mouse subcutaneous injection model. (A) Images of representative lesion-bearing mice injected with the indicated C2C12 clones (top text, M# indicates mouse number in the group) at the time of sacrifice (bottom text). Red arrows indicate lesion sites.
